# How to make more from exposure data? An integrated machine learning pipeline to predict pathogen exposure

**DOI:** 10.1101/569012

**Authors:** Nicholas M. Fountain-Jones, Gustavo Machado, Scott Carver, Craig Packer, Mariana Recamonde-Mendoza, Meggan E. Craft

**Author notes:** Department of Veterinary Population Medicine, University of Minnesota, 1365 Gortner Avenue, St Paul, Minnesota 55108 (Nicholas M. Fountain-Jones). **Data Accessibility Statement** Should the manuscript be accepted, the data supporting the results will be archived on GitHub and Dryad with DOI provided.

## Abstract

1. Predicting infectious disease dynamics is a central challenge in disease ecology. Models that can assess which individuals are most at risk of being exposed to a pathogen not only provide valuable insights into disease transmission and dynamics but can also guide management interventions. Constructing such models for wild animal populations, however, is particularly challenging; often only serological data is available on a subset of individuals and non-linear relationships between variables are common.
2. Here we take advantage of the latest advances in statistical machine learning to construct pathogen-risk models that automatically incorporate complex non-linear relationships with minimal statistical assumptions from ecological data with missing values. Our approach compares multiple machine learning algorithms in a unified environment to find the model with the best predictive performance and uses game theory to better interpret results. We apply this framework on two major pathogens that infect African lions: canine distemper virus (CDV) and feline parvovirus.
3. Our modelling approach provided enhanced predictive performance compared to more traditional approaches, as well as new insights into disease risks in a wild population. We were able to efficiently capture and visualise strong non-linear patterns, as well as model complex interactions between variables in shaping exposure risk from CDV and feline parvovirus. For example, we found that lions were more likely to be exposed to CDV at a young age but only in low rainfall years.
4. When combined with our data calibration approach, our framework helped us to answer questions about risk of pathogen exposure which are difficult to address with previous methods. Our framework not only has the potential to aid in predicting disease risk in animal populations, but also can be used to build robust predictive models suitable for other ecological applications such as modelling species distribution or diversity patterns.

## Introduction

An individual’s risk of infection by a pathogen is dependent upon a wide variety of host and pathogen traits that are often complicated by heterogeneities in landscapes and individual behavior (Altizer et al., 2003; Carver et al., 2016; Ezenwa, 2004; Nunn, Jordán, McCabe, Verdolin, & Fewell, 2015). Evidence of infection in animals is often determined by serology, which can determine if a host has likely been infected (or not) sometime in the past (‘exposed’ vs ‘non-exposed’ host, Gilbert et al 2013). Models that attempt to quantify exposure risk often assume exposed and non-exposed hosts can be linearly separable by a predictor even though non-linear relationships may be more biologically plausible, e.g., where exposure risks reach an asymptote by a certain age (Denison & Holmes, 2001). Further, exposure risk may be dependent on interactions between predictors (Diez-Roux, 1998) (e.g., exposure risk is only elevated in the youngest and oldest individuals in one subset of the population). Generalized linear models are not effective in capturing such non-linear responses and complex interactions, which is an important advantage of most machine-learning algorithms (Elith, Leathwick, & Hastie, 2008; Tu, 1996).

Machine learning methods offer a powerful and flexible alternative to explore important patterns in exposure data, often with substantially better predictive performance (e.g., Elith et al., 2008). Machine learning models can include thousands of predictors without overfitting and can analyze either categorical or continuous data, even in datasets containing large numbers of missing values. However, choosing the most appropriate machine learning model can be challenging, thus the development of methods for identifying the best-performing model has become an important area of research over the last 15 years (Ho & Pepyne, 2002). Supervised machine learning models, such as support vector machines (SVMs), or tree-based algorithms, such as random forests (RF), gradient boosting models (GBMs), or boosted regression trees (BRTs), operate in markedly different ways, potentially leading to different performance and results (Machado, Mendoza, & Corbellini, 2015). While machine learning models are increasingly popular in disease ecology and epidemiology (e.g., Babayan, Orton, & Streicker, 2018; Han et al., 2016; Hollings et al., 2017), they have not yet been widely adopted.

In part, the limited application of machine learning models to disease ecology is due to the perception of these models as black boxes. Recent advances have helped unveil the black box by clarifying the predictive capabilities of these models (Brdar, Gavrić, Ćulibrk, & Crnojević, 2016; Lundberg & Lee, 2016; Molnar, 2018b). For example, coalitional game theory can be used to untangle how variables contribute to a machine learning model prediction (Štrumbelj & Kononenko, 2014), providing an explanation for why an individual host was predicted to be positive or negative for pathogen exposure.

The conditions of when and where an individual was infected by a pathogen are often unknown (Gilbert et al., 2013). Exposure risk is often quantified by analyzing variables such as age or group size or landscape context based on the date the individual was sampled, even though the start of the infection was likely to have occurred sometime in the past (Carver et al., 2016; Packer et al., 1999). Further, risk factors of pathogen exposure are often assumed to be fixed in time, even though the ecological context could change between exposure and time of sampling (Goldstein et al., 2017; Munson et al., 2008). The spatio-temporal mismatch between the location and date of infection and the location and date of sampling is well known (Gilbert et al., 2013; Goldstein et al., 2017), yet few pathogen risk models have explicitly considered these limitations.

Here we introduce a multi-algorithm machine learning ensemble pipeline that incorporates recent advances in data science to better predict pathogen exposure risk. Our pipeline is flexible and many of the post-processing methods are model agnostic in that they can be used to interrogate any regression or classification model. To demonstrate the utility of our approach, we model pathogen exposure risk in two different viruses that infect African lions (*Panthera leo*): canine distemper virus (CDV) and feline parvovirus (parvovirus). To build our exposure risk models, we leverage age estimates, long term behavioural observations, and environmental data from the Serengeti Lion Project (SLP) as predictors (hereafter called ‘features’ in line with computer science terminology). Using this data, we develop exposure risk models based on features calibrated to reflect potential date of exposure (rather that sampling date), and also compare exposure risk factors between the two pathogens. While we focus on pathogens in wildlife populations, our pipeline could be applied in a similar way to human, domestic animal, or livestock exposure data, as well as to any classification or regression problem. We provide code and a fully worked example in a vignette (Text S1).

## Material and Methods

If exposure data has been collected across multiple years and there are reliable estimates of host age, researchers can take advantage of our calibration steps below. If not, researchers can proceed to *2.2 Pre-processing*.

### 2.1 Data calibration

Plotting prevalence relationships across host age and time is an essential first step to start calibrating exposure models. When evaluating year-prevalence curves, spikes in prevalence in certain years may estimate the timing of likely epidemics, particularly when the year-prevalence curves of juveniles are considered (Fig. 1)(Packer et al., 1999). Knowledge of pathogen natural history (e.g., chronic vs acute infection) and tests such as chi-squared testing for temporal fluctuations can be used to support these estimates (Fig. 1., Packer et al., 1999). In populations with more precise age estimates beyond juvenile/adult classes, age of likely exposure can be calculated based on how old the individual was when the last epidemic occurred. All other features can be calibrated to the year of exposure (e.g., what was the group size for that individual for the potential year of exposure). Some host characteristics such as sex do not change with time. If the analysis uses serological evidence, detection of exposure is often based on a cut-off threshold (i.e., above a certain titre an individual is classified as exposed). This is imperfect as antibody titre tends to diminish over time (Gilbert et al., 2013; Packer et al., 1999), therefore adding ‘time since epidemic’ as a feature can account for imperfect detection. Accounting for imperfect detection can allow for more accurate measures of infection risk in disease surveys (Lachish, Gopalaswamy, Knowles, & Sheldon, 2012)

Age-prevalence relationships can also help calibrate possible exposure times for pathogens more endemic, or constantly present, in a population (Fig. 1). If prevalence peaks at an early age and is stable over time, this may be an indication that the pathogen is more endemic in the population; the relevant features can be calibrated to reflect that the individual was likely exposed early in life (Fig. 1). Age-specific force of infection models (FOI, or the per capita rate at which susceptible individuals acquire infection) can help support these estimates (i.e., at what age is the FOI highest; Anderson & May, 1985). If prevalence increases in a linear fashion with age and the FOI is equal across age groups (Fig. 1), gaining a reliable estimate of the year of exposure (and therefore calibration, as described below) is difficult.

Depending on the interpretation of the age- and year-prevalence curves, temporal features can be estimated, or calibrated, including age exposed, time since last epidemic, last epidemic year, year exposed, and age sampled (Fig. 1).

**Fig.1:**
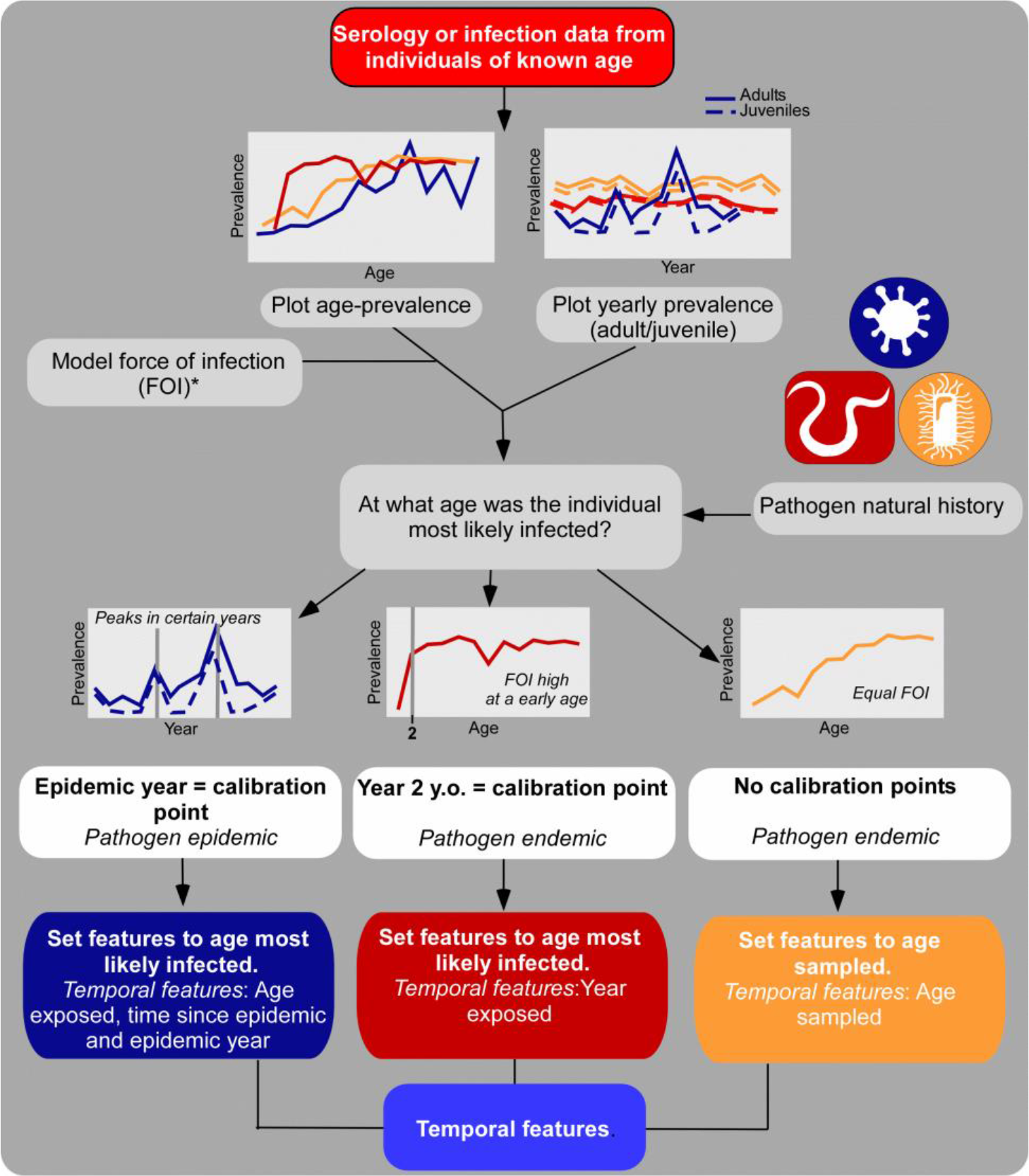
Flow chart showing the pre-processing steps in our machine learning pipeline. Each colour represents a separate pathogen. FOI: Force of infection. *: indicates optional step.

### 2.2 Pre-processing

It is important to account for missing data either by imputation or removal prior to model construction (Fig. 2). Some machine learning algorithms, such as gradient boosting, bin missing data as a separate node in the decision tree (Friedman, 2002, Fig. S1), however other algorithms such as SVM are less flexible. In order to compare predictive performance across models, missing data can either be imputed or removed from the dataset. Although providing specific advice on whether to include missing data or not is outside the scope of this paper (see Nakagawa & Freckleton, 2008), we provide an option if imputation is suitable for the study problem. We integrated the ‘missForest’(Stekhoven & Bühlmann, 2012) machine-learning imputation routine (using the RF algorithm) into our pipeline, as it has been found to have low error rates with ecological data (Penone et al., 2014).

**Fig. 2:**
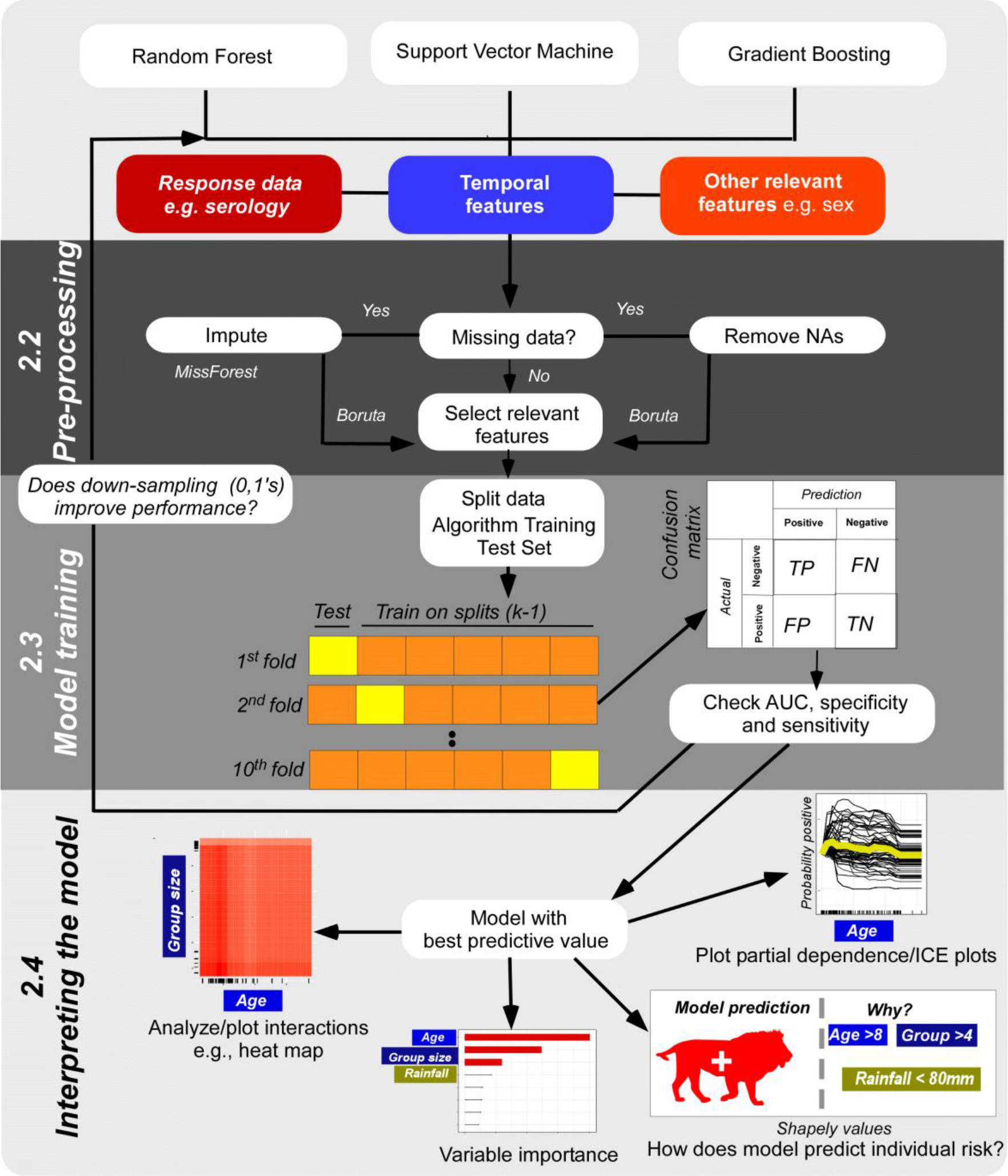
Flow chart showing the steps in our machine learning pipeline. Vertical text in bold indicates how the chart relates to sections of the main text and vignette (Text S1, sections 1-3 respectively). TP: True positive, FP: False positive, FN: False negative, TN: True negative. Yellow boxes indicate which data split is being tested in that particular ‘fold’.

Another important step before constructing a model, particularly for relatively small data sets with large numbers of features, is to select features that are relevant and informative for prediction. Reducing the feature set used in models not only increases the efficiency of the algorithms, but also improves the accuracy of many machine learning algorithms (Kohavi & John, 1997; Kursa & Rudnicki, 2010). In our pipeline, we use the ‘Boruta’ (Kursa & Rudnicki, 2010) feature selection algorithm for reducing the feature set to just those relevant for prediction prior to model building. The Boruta routine has been found to be the one of the most powerful approaches to select relevant features (Degenhardt, Seifert, & Szymczak, 2017) and does so by measuring the importance of each variable using an RF-based algorithm (Kursa & Rudnicki, 2010).

### 2.3 Model training

We incorporated an internal repeated 10-fold cross-validation (CV) process to estimate model performance. CV can help prevent overfitting and artificial inflation of accuracy due to use of the same data for training and validation steps (Fig. 2). To run and evaluate each model, our pipeline uses the ‘caret’ (classification and regression training) package in R (Kuhn, 2008). Not only does this package provide a streamlined approach to tuning parameters for a wide variety of models, including machine learning models, it also offers functions that directly enable robust comparisons of model performance. The ‘train’ function uses resampling to evaluate how tuning parameters such as learning rate (see Elith et al., 2008) can impact model performance and ‘chooses’ the optimal model with the highest performance via the confusion matrix (e.g., sensitivity and specificity for classification models). Another advantage of this package is that it can perform classification or regression using 237 different types of models from generalized linear models (GLMs such as logistic regression) to complex machine learning and Bayesian models using a standardized approach (see Kuhn, 2008 for a complete list of models).

In our pipeline, we compare supervised machine learning algorithms (RF, SVM, and GBM) as well as logistic regression. These models are among the most popular and best tested machine learning methods, but all operate in different ways, and this can in turn can impact predictive performance as measured by AUC (Marmion, et al., 2009; Ogutu, Piepho, & Schulz-Streeck, 2011). In brief, for classification problems, SVMs use vectors and kernel functions that maximizes the margin between classes of data on a hyperplane (see Schölkopf & Smola, 2002 for model details). In contrast, RF and GBM are tree-based algorithms that iteratively split data into increasingly pure ‘sets’, with the main difference being that GBM fits trees sequentially with each new tree helping correct errors from the previous (Friedman, 2002). For RF, the trees are fit independently with each iteration (see Fig. S1 for a more detailed comparison).

Imbalanced proportions of the outcome of interest are common in disease ecology, where for example, prevalence of the pathogen is < 50%; imbalanced proportions can influence the predictive performance of the algorithm (Haibo He & Garcia, 2009). To address this issue, we included a down-sampling strategy in our pipeline that can be used to compare model performance for datasets with imbalanced data (Fig. 2).

### 2.4 Interpreting the model

Once the best model is selected, we provide a variety of tool sets to visualize and interrogate model results (Fig. 2) based on the ‘iml’ (interpretable machine learning) R package (Molnar, 2018a). Measures of variable importance are often used to understand which features contributed to the model in ranked order of importance. However, comparing variable importance across multiple types of models can be challenging as models often quantify variable importance in different ways. Further, in some cases, different models can have the same predictive performance but rely on different features or the ‘Rashomon’ effect (Breiman, 2001). Here, to help overcome these limitations, our pipeline presents a model agnostic method called ‘model class reliance’ (Fisher, Rudin, & Dominici, 2018) as an extension of Breiman’s (2001) permutation approach. In essence, the algorithm quantifies the expected loss in predictive performance (i.e., how well the model classifies exposed vs non-exposed individuals) for a pair of observations compared to the full model in which the particular feature has been switched (see Fisher et al., 2018; Molnar, 2018b for more details). Because this algorithm compares the ratio of error rather than the differences in model error rates it can be used to compare different models. A feature is not considered important if switching it does not alter model performance. Further, this method is probabilistic and has the advantage that feature permutation automatically includes feature interaction effects into account into the importance calculation (Molnar, 2018b).

Our pipeline also provides the appropriate code to calculate and plot both partial dependence plots (PD) and individual conditional expectation (ICE) to examine the marginal effect of a particular feature on the response (Goldstein et. al., 2015). PD plots visualize the global (i.e., average model) effect of a feature on the response, whilst ICE plots examine the effect of each feature for each observation, whilst holding all other features constant (Goldstein et al., 2015; Molnar, 2018b). For example, to assess the effect of a feature on exposure risk of individual ‘A’, the ICE approach would vary the feature across a grid of values. For each feature variant, the selected machine learning model is used to make new predictions of exposure risk for individual A. Then a line is fit to the predictions across feature values and then the process is repeated for each individual observation. The PD plot represents the average of the lines an ICE plot (e.g., yellow line is the average whereas the black lines are the individual observations in the bottom panel of Fig. 2). Combined they overcome limitations of each approach separately as PD plots alone obscure when interactions shape a feature-response relationship, whilst ICE plots can be challenging to read without plotting the average effect (Molnar, 2018b). We recommend ICE plots centred on the minimum feature value (cICE) in order to more easily visualize when ICE estimates vary between observations (Molnar, 2018a).

To further interrogate interactions in each model, our pipeline provides the option to calculate Friedman’s H-statistic (Friedman & Popescu, 2008) for all features (and combinations thereof) in the model. The strongest interactions can then be visualized using 3D PD plots or heatmaps (Fig. 2). In brief, Friedman’s H-statistic is a flexible way to quantify feature interaction strength using partial dependency decomposition and represents the portion of variance explained by the interaction (see Friedman & Popescu, 2008 for details). Whilst computationally expensive, this method is also model agnostic and can be upscaled to quantify high order interactions, i.e., three- or four-way interactions between features (Molnar, 2018b).

Finally, our pipeline provides functions to calculate Shapely values (Shapley, 1953) to assess how individual features effect the prediction of single observations (i.e., an individual lion). This approach calculates Shapely values from the final model that assign ‘payouts’ to ‘players’ depending on their contribution towards the prediction (Brdar et al., 2016; Molnar, 2018b; Štrumbelj & Kononenko, 2014). The players cooperate with each other and obtain a certain ‘gain’ from that cooperation. In this case, the ‘payout’ is the model prediction, ‘players’ are the feature values of the observation (e.g., host sex) and the ‘gain’ is the model prediction for this observation (e.g., was the host exposed or not) minus the average prediction of all observations (on average what is probability of a host being exposed) (Molnar, 2018b). More specifically, Shapely values, defined by a value function (*Va*), compute feature effects (*ϕ*_*ij*_) for any predictive model *f*^(*i*)^:

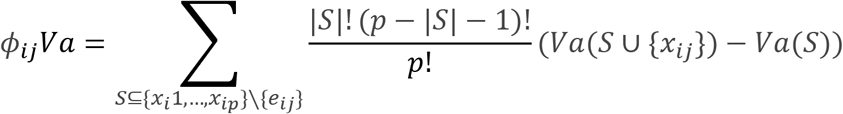

Where *S* is a subset of the features (or players), *x* is the vector features values for observation *i* and *p* is the number of features. *Va*_*xi*_ is the prediction of feature values in *S* marginalized by features not in *S*:

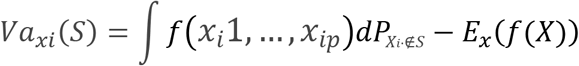

### 2.5 Modelling lion exposure risk

Blood samples were collected from 300 individual lions of known age (estimated to the month) in the Serengeti Ecosystem, Tanzania between 1984 and 1994. Of these individuals, 40% were seropositive for CDV and 30% for parvovirus (see Hofmann-Lehmann et al., 1996; Roelke-Parker et al., 1996 for method details). Both viruses can infect multiple species, yet CDV is directly transmitted via aerosol, whereas parvovirus can be either transmitted directly or through environmental contamination. CDV and parvovirus are considered to have epidemic cycles in the Serengeti lion population (Hofmann-Lehmann et al., 1996; Packer et al., 1999). Chi-square tests of year-prevalence relationships supported three epidemic years for both viruses: ~1977, 1981 and 1994 for CDV and 1977, 1985 and 1992 for parvovirus (Packer et al., 1999). Subsequent Bayesian state-space analysis of this data further confirmed these years as epidemic years for CDV and parvovirus (Behdenna et al., *in press*; Viana et al., 2015).

We selected 17 features and calibrated them where appropriate (as detailed in *2.1 Data calibration*). See Text S2 for feature selection details. Each model, including logistic regression, was performed following the steps outlined in our pipeline. We compared the predictive performance of the models using calibrated and uncalibrated feature sets (hereafter calibrated or uncalibrated models) to assess how differences in calibration could change the exposure risk predictions. Uncalibrated feature sets were calculated based on the date an individual was sampled, rather than going through the process outlined in *2.1 Data calibration*.

## Results

### Machine learning models with calibrated feauture sets have higher predictive performance

For both CDV and parvovirus, machine learning models had higher predictive performance (higher AUC) compared to logistic regression models (Table 1). For example, predictions of parvovirus were only just better than random using logistic regression model (AUC = 0.54), whereas the GBM model was correctly predicting exposure 73% of the time (AUC = 72.98). Using our calibration approach further improved the overall predictive performance of each pathogen by increasing the sensitivity of the models (i.e., they more able to correctly identify positives), however, there was a trade-off with reduced specificity. For example, our calibrated CDV model had a 7% increase in AUC with an 23% increase in sensitivity but 18% decrease in specificity compared to the uncalibrated model (Table 1). The features considered relevant in each model were relatively consistent (Table 1). However, there were some important exceptions with, for example, sex only relevant for parvovirus exposure in the calibrated model. In contrast, rainfall was relevant for predicting parvovirus exposure in the uncalibrated model but not in the calibrated model.

**Table 1:**
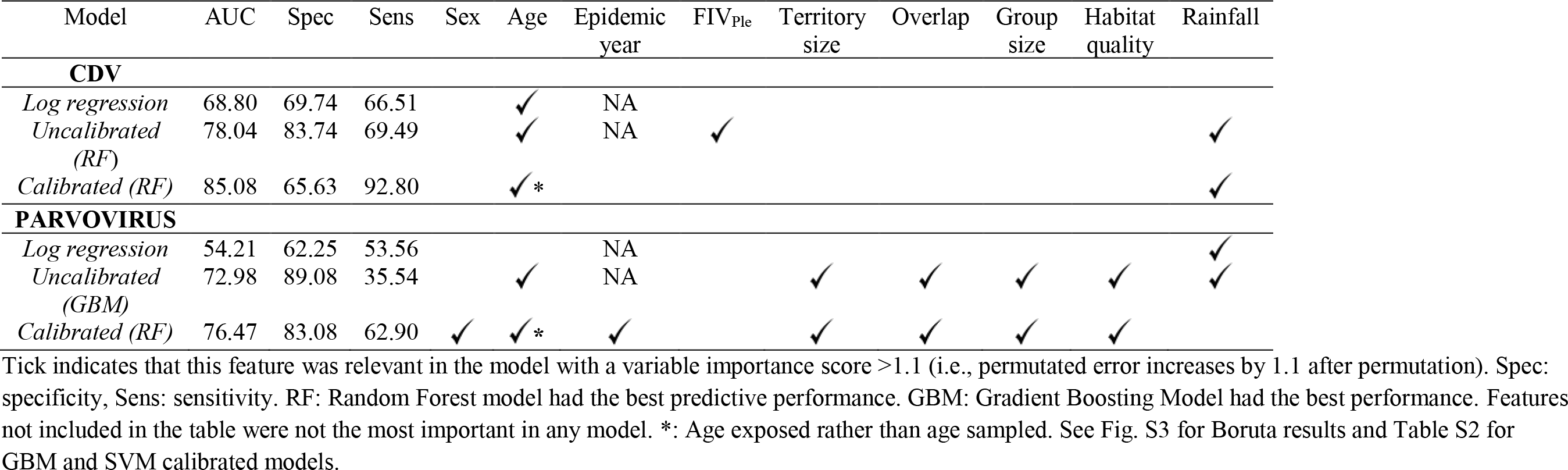
Model performance and relevant features for each model. If a feature was not relevant in any model it was excluded using the Boruta algorithm.

### Model calibration alters feature-risk relationships

Age and rainfall were the most important features predicting CDV exposure, but both features were relatively less important in the calibrated models (Fig. 3). Even though the features associated with exposure risk in each model were broadly similar for both pathogens, the relationships between each feature and exposure risk varied. Partial dependency plots showed that risk of CDV increased relatively linearly across age classes in in the uncalibrated model (Fig. 3b), whereas in the calibrated model exposure risk was much more constant across age classes with an increase in risk in individuals between 1-2 y.o. (Fig. 3f). Rainfall also showed different relationships in each model with reduced exposure risk when the average monthly rainfall > 40 mm in the age calibrated model (Fig. 3c). There was a much shallower decline in CDV risk associated with rainfall in the calibrated model (Fig 3e) compared to the uncalibrated model (Fig. 3c).

**Fig. 3:**
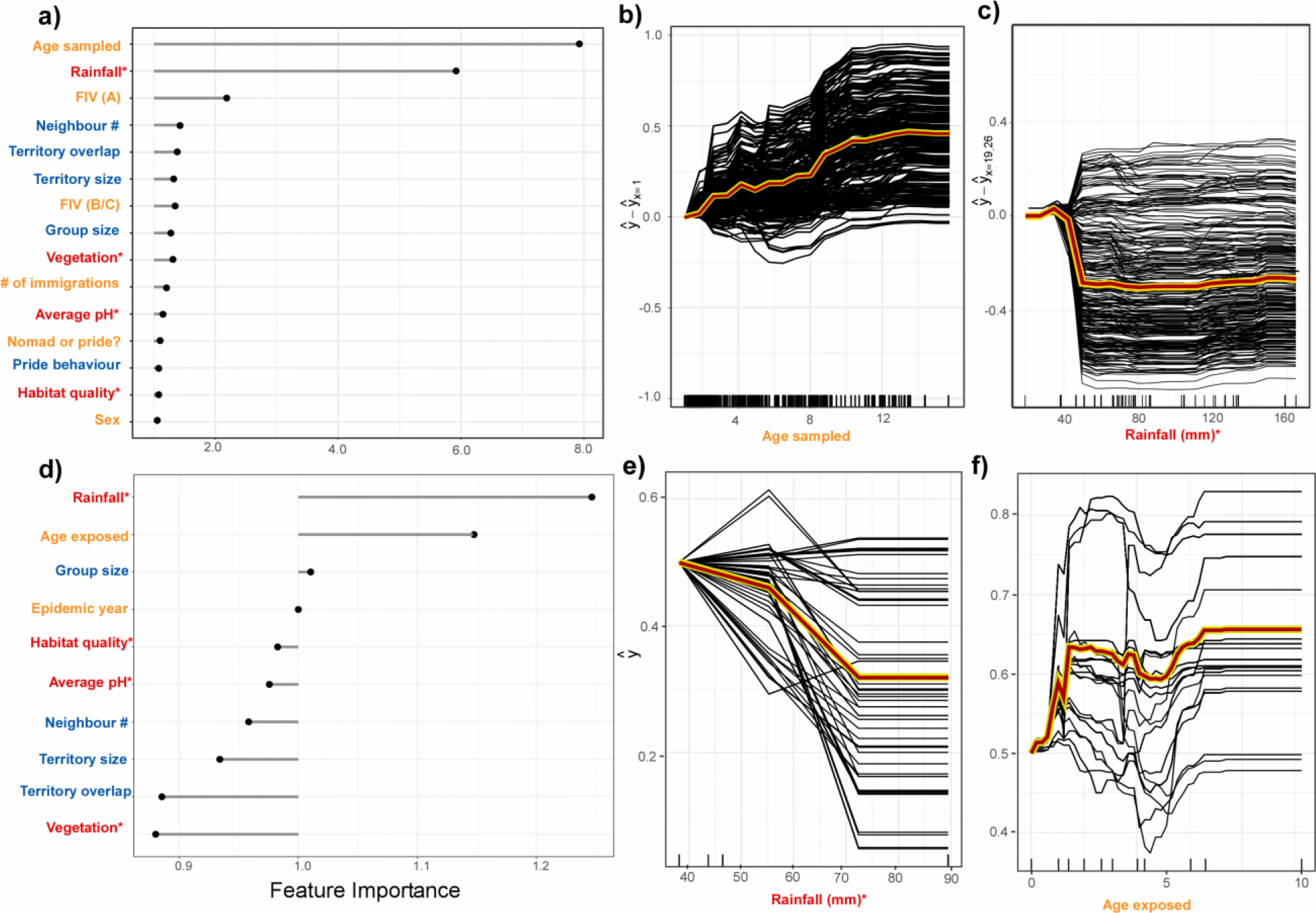
Plots showing the differences in model predictions and the features that contribute to CDV exposure risk in the Serengeti lions for the uncalibrated models (a-c) compared to the calibrated models (d-f). Age sampled followed by rainfall are the most important features associated with CDV exposure risk in the uncalibrated models (a). In contrast, rainfall followed by age exposed were the most important features in the calibrated model (d). However, cICE plots show that the relationship with exposure risk differs substantially (b & c: uncalibrated model, e & f: calibrated model). Feature name colours reflect feature type (orange = individual, blue = pride-level, red = environmental) and * indicates that the variable was averaged across the pride’s territory. See Text S2 for further details including sample size. Tick marks on the x-axes (b, c, e & f) show the distribution of data. Yellow/red line indicates the average value across all individuals. The y-axes reflect probability of being positive for CDV.

Similar to CDV, age sampled followed by rainfall were the most important features associated with parvovirus exposure risk in the uncalibrated models (Fig. S4a). Parvovirus exposure risk slightly increased across age classes in the uncalibrated models, however in the calibrated models exposure risk increased rapidly at early ages (0-1), but then was relatively constant across ages >3 (Fig. S4b). The signature of rainfall on parvovirus risk in the uncalibrated model was remarkably like that of CDV with a large drop in risk when the monthly rainfall was > 40 mm a month (Fig. S4c). However, rainfall was much less important in the calibrated model (Fig. S4d). Strikingly epidemic year was important in the calibrated model with exposure risk much higher for animals likely exposed in the 1992 epidemic (Fig. S4f).

### Interactions

We further interrogated the calibrated models to visualize how interactions between features could be important for exposure risk of both pathogens. We focussed on interactions with epidemic year, as we were interested to see if exposure risk could vary with each outbreak (see Fig. S5 for a summary of all interactions detected). For CDV, the strongest interaction with epidemic year was age exposed (Fig. 4a). Even though exposure risk was predicted to be reasonably similar in each CDV outbreak (Fig. S6), exposure risk was higher for lions 2-5 y.o. in the 1994 epidemic compared to the 1981 epidemic (Fig. 4b). For parvovirus, territory overlap had the strongest interaction with epidemic year (Fig. 4c) with individuals in prides with low overlap having an increased risk of parvovirus during the 1992 epidemic only. There was also an interaction with epidemic year and age exposed with lions aged 10-12 y.o. more likely to be exposed to parvovirus during the 1992 epidemic (Fig. 4d). See Text S3 and Fig. S7 for a description of how individual features effects the prediction of individual lion’s exposure risk.

**Fig. 4:**
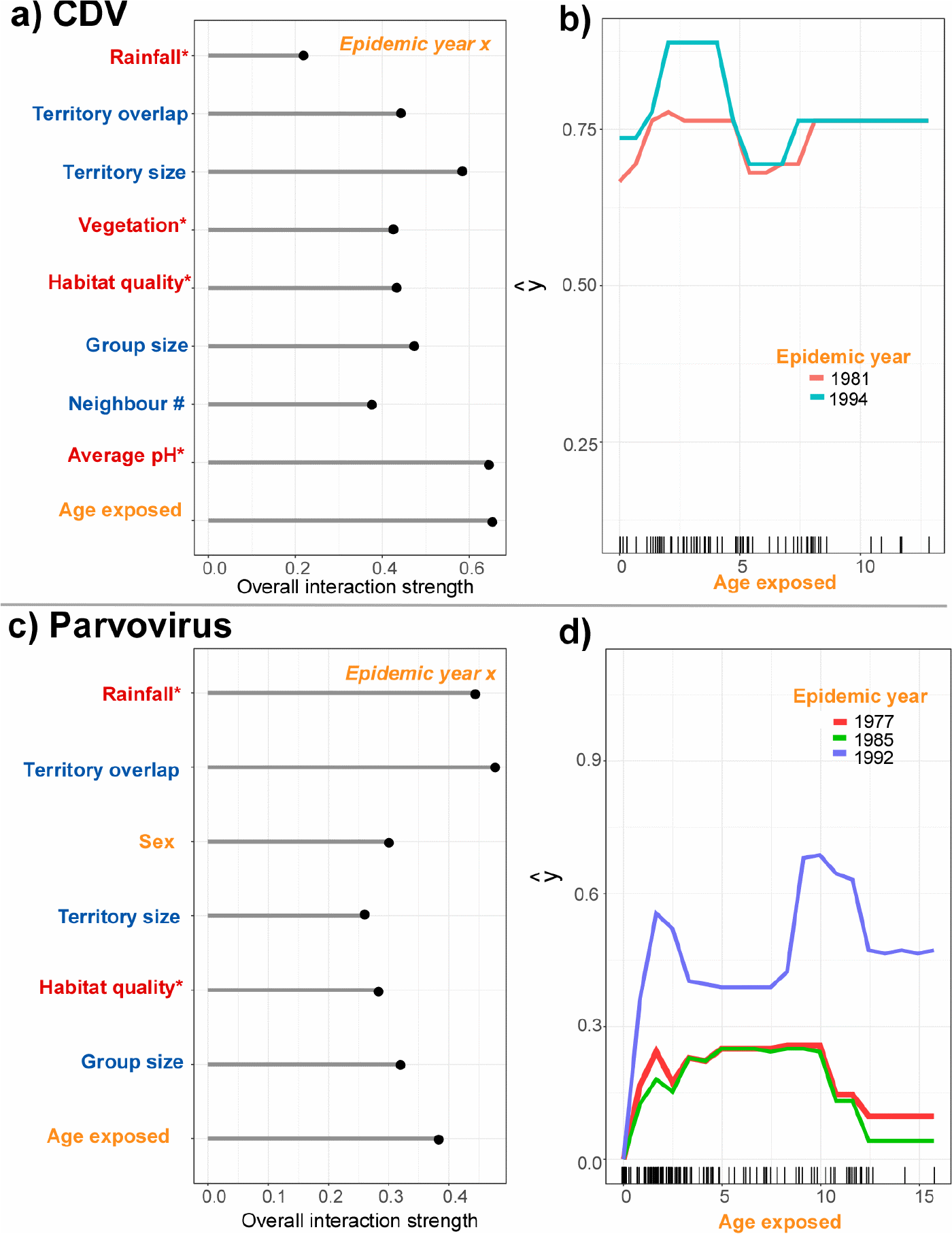
Interactions between epidemic year and the other features in shaping exposure risk of CDV (a-b) and parvovirus (c-d) in the Serengeti lions. Differing coloured partial dependence curves (b & d) represent different epidemics. The y axis (y hat) is the relative risk of being exposed (with all other feature combinations marginalized). See Fig. S5 for the interactions detected for other features.

## Discussion

Our machine learning pipeline offers a powerful approach that can enable researchers to generate robust exposure risk models and interrogate models in new ways. We found that models, where host and environmental features were calibrated based on likely exposure year, had higher predictive power compared to uncalibrated models (i.e., comprised of features based on sample date). The calibrated models not only provided better predictive power but also more robust insights into pathogen dynamics in a wild population that can aid future surveillance. Furthermore, use of this pipeline is not restricted to research on pathogen exposure, but also could be employed with minimal change to model other ecological systems.

### Robust insights in exposure risk for CDV and parvovirus

The machine learning algorithms employed in our pipeline had higher predictive performance compared to the logistic regression models because they were able to quantify non-linear relationships in our data. For both pathogens, non-linear relationships between host and environmental features were common and provided new perspectives of pathogen dynamics in this lion population. For example, in our calibrated model of parvovirus, risk was saturated when individuals > 3 y.o but decreased after age 7 (Fig. S4d-e). Puppies are thought to be the highest risk category for canine parvovirus (Pollock & Coyne, 1993) and, based on our calibrated models, this may be true for feline parvovirus in lions as well. Even though this pattern is relatively intuitive, neither the logistic regression (Table 1), the uncalibrated model (Fig. S4a-b), or age-prevalence curves (Fountain-Jones et al., *in press*) captured this pattern, highlighting the need for these types of models to quantify exposure risk.

Calibrating features provided not only a more nuanced interpretation of pathogen dynamics but also higher predictive performance. The greater predictive performance could be because we removed individuals that were not on the landscape during the inter-epidemic years, and thus were negative for the pathogen, not due a behavioral or landscape trait, but due to the cyclic nature of epidemics. As exposure risk changed over time, including ‘epidemic year’ in the models may have been responsible for the greater predictive power of the calibrated models. Further, even though it was not relevant in the best performing CDV and parvovirus models, including ‘time since exposure’ as a feature provides support that the observed relationships were less likely to be due to differences in detectability due to titre decline over time(Packer et al., 1999).

Another advantage of the machine learning methods is the ability to model the non-linear interactions shaping exposure risk in the lions. In our case we focused on how host and environmental characteristics interacted with epidemic year to better understand the temporal dimension of exposure risk: an important question rarely tested in wildlife populations (Gilbert et al., 2013). For parvovirus, in particular, the risk associated with lion age varied between epidemics with the 1992 epidemic standing out with increased risk in older as well as younger individuals (Fig. 4). For CDV in contrast, there were remarkably similar risk factors for the 1981 and 1994 epidemics, except for an increase in risk in juveniles in 1994 (Fig. 4b). However, there was no known increase in mortality in 1981 as there was in the 1994, further supporting the idea that co-infection with *Babesia* was the principal agent driving mortality (Munson et al., 2008). When and why exposure risk is stable or dynamic across epidemic years is an open question in disease ecology, and analysis pipelines such as ours can help address this gap.

Further, calculating Shapely values from predictive models can unpack what each model means in terms of exposure risk in an intuitive way that can be used to inform practitioners and guide surveillance and forecast future risk. For example, when rainfall is below average individuals from prides with relatively high overlap are more at risk of being exposed to CDV perhaps due to increased chances of congregating at waterholes. The sex and age of individuals are less important when choosing which individuals to test or vaccinate.

### Caveats and future directions

Even though serological or qPCR evidence is often used (including here) as evidence of past or current infection, there are numerous limitations associated with this type of data that need to be considered (Gilbert et al., 2013; Lachish et al., 2012). Further, for pathogens that cause high morality, the individuals sampled represent survivors of the epidemic, and this may bias exposure risk models for wildlife populations that are infrequently monitored. For example, if males experienced high mortality in a particular epidemic compared to females, post epidemic sampling of this population may incorrectly infer that females were more likely to be exposed. How to incorporate biases such as these into exposure risk models could be a fruitful future direction for research.

A more general limitation of this approach is that uncertainty in model predictions for the machine learning algorithms we used is not well characterized. Bayesian additive regression trees (BART) may be a promising technique that could fill this gap, but this technique comes at a computational cost that may be prohibitive for larger datasets (Chipman, George, & McCulloch, 2010). Furthermore, none of the methods we employed in our pipeline explicitly account for spatial relationships which could be a limitation for studies that use data across a larger geographic area where spatial autocorrelation could be an important problem.

Lastly this analysis pipeline could be used for a variety of ecological applications including both classification and regression problems with continuous or categorical response variables. For example, our pipeline could easily be incorporated into species distribution models. Moreover, as the R package ‘caret’ is highly flexible and can perform and compare a large number of algorithms with minimal change to the R syntax, the end user could easily test any model of their choosing in a similar way (e.g. generalized additive models or discriminant analysis). Similarly, the code we provide from the R package ‘iml’ is model agnostic and can provide an increase in the interpretability for many regression or classification models (Molnar, 2018a).

### Conclusions

Our analysis pipeline offers a flexible unified approach to not only better understand pathogen exposure risk, but also could be applied to any system where non-linear responses and interactions are important in shaping a response variable. This study has highlighted the value of using advances in data science coupled with feature calibration to increase our understanding of how host and landscape characteristics shape pathogen risk in populations, and how exposure risk could change over time. Machine learning approaches are rapidly evolving and future developments coupled with advances in pathogen detection are likely to provide even more resolution on the drivers of pathogen exposure risk.

## Acknowledgements

N.M. F-J. and M.E.C. were funded by National Science Foundation (DEB-1413925) and M.E.C. was funded by National Science Foundation (DEB-1654609) and CVM Research Office UMN Ag Experiment Station General Ag Research Funds.

**Author Contributions**

The study was designed by NMFJ, CP and MEC and the data were provided by CP. NMFJ, GM and MRM conducted all statistical analyses, MRM led the development of the R optimization functions, and NMFJ wrote the first draft of the manuscript. All authors (NMFJ, GM, SC, CP, MRM and MEC) contributed substantially to revisions. NMFJ and GM prepared the vignette.

## References

Altizer, S., Nunn, C. L., Thrall, P. H., Gittleman, J. L., Antonovics, J., Cunningham, A. a., … Pulliam, J. R. C. (2003). Social organization and parasite risk in mammals : Integrating theory and empirical studies. Annual Review of Ecology, Evolution, and Systematics, 34(1), 517–547. doi:10.1146/annurev.ecolsys.34.030102.151725

Anderson, R. M., & May, R. M. (1985). Age-related changes in the rate of disease transmission: implications for the design of vaccination programmes. The Journal of Hygiene, 94(3), 365–436. Retrieved from http://www.ncbi.nlm.nih.gov/pubmed/4008922

Babayan, S. A., Orton, R. J., & Streicker, D. G. (2018). Predicting reservoir hosts and arthropod vectors from evolutionary signatures in RNA virus genomes. Science, 362(6414), 577–580. doi:10.1126/science.aap9072

Behdenna, A., Lembo, T., Cleaveland, S., Halliday, J. E. B., Packer, C., Lankester, F., … Viana, M. (in press). Transmission ecology of canine parvovirus in a multi-host, multi-pathogen system. Proc R Soc B.

Brdar, S., Gavrić, K., Ćulibrk, D., & Crnojević, V. (2016). Unveiling spatial epidemiology of HIV with mobile phone data. Scientific Reports, 6(1), 19342. doi:10.1038/srep19342

Breiman, L. (2001). Random forests. Machine Learning, 45(1), 5–32. doi:10.1023/A:1010933404324

Carver, S., Bevins, S. N., Lappin, M. R., Boydston, E. E., Lyren, L. M., Alldredge, M., … VandeWoude, S. (2016). Pathogen exposure varies widely among sympatric populations of wild and domestic felids across the United States. Ecological Applications, 26(2), 367–381.

Chipman, H. A., George, E. I., & McCulloch, R. E. (2010). BART: Bayesian additive regression trees. The Annals of Applied Statistics, 4(1), 266–298. doi:10.1214/09-AOAS285

Denison, D. G. T., & Holmes, C. C. (2001). Bayesian partitioning for estimating disease risk. Biometrics, 57(1), 143–149. doi:10.1111/j.0006-341X.2001.00143.x

Diez-Roux, A. V. (1998). Bringing context back into epidemiology: variables and fallacies in multilevel analysis. American Journal of Public Health, 88(2), 216–22. doi:10.2105/AJPH.88.2.216

Elith, J., Leathwick, J. R., & Hastie, T. (2008). A working guide to boosted regression trees. Journal of Animal Ecology, 77(4), 802–813. doi:10.1111/j.1365-2656.2008.01390.x

Ezenwa, V. O. (2004). Host social behavior and parasitic infection: a multifactorial approach. Behavioral Ecology, 15(3), 446–454. doi:10.1093/beheco/arh028

Fisher, A., Rudin, C., & Dominici, F. (2018). Model class reliance: Variable importance measures for any machine learning model class, from the “Rashomon” Perspective. Retrieved from http://arxiv.org/abs/1801.01489

Fountain-Jones, N.., Packer, C., Jacquot, M., Blanchet, G., Terio, K., & Craft, M.. (in press). Endemic infection can shape exposure to novel pathogens: Pathogen co-occurrence networks in the Serengeti lions. Ecology Letters.

Friedman, J. H. (2002). Stochastic gradient boosting. Computational Statistics & Data Analysis, 38(4), 367–378. doi:10.1016/S0167-9473(01)00065-2

Friedman, J. H., & Popescu, B. E. (2008). Predictive learning via rule ensembles. The Annals of Applied Statistics, 2(3), 916–954. doi:10.1214/07-AOAS148

Gilbert, A. T., Fooks, A. R., Hayman, D. T. S., Horton, D. L., Müller, T., Plowright, R., … Rupprecht, C. E. (2013). Deciphering serology to understand the ecology of infectious diseases in wildlife. EcoHealth, 10(3), 298–313. doi:10.1007/s10393-013-0856-0

Goldstein, A., Kapelner, A., Bleich, J., & Pitkin, E. (2015). Peeking inside the black box: Visualizing statistical learning with plots of individual conditional expectation. Journal of Computational and Graphical Statistics, 24(1), 44–65. doi:10.1080/10618600.2014.907095

Goldstein, E., Pitzer, V. E., O’Hagan, J. J., & Lipsitch, M. (2017). Temporally varying relative risks for infectious diseases. Epidemiology. NIH Public Access. doi:10.1097/EDE.0000000000000571

Han, B. A., Schmidt, J. P., Alexander, L. W., Bowden, S. E., Hayman, D. T. S., & Drake, J. M. (2016). Undiscovered bat hosts of Filoviruses. PLOS Neglected Tropical Diseases, 10(7), e0004815. doi:10.1371/journal.pntd.0004815

Ho, Y. C., & Pepyne, D. L. (2002). Simple explanation of the no-free-lunch theorem and its implications. Journal of Optimization Theory and Applications, 115(3), 549–570. doi:10.1023/A:1021251113462

Hofmann-Lehmann, R., Fehr, D., Grob, M., Elgizoli, M., Packer, C., Martenson, J. S., … Lutz, H. (1996). Prevalence of antibodies to feline parvovirus, calicivirus, herpesvirus, coronavirus, and immunodeficiency virus and of feline leukemia virus antigen and the interrelationship of these viral infections in free-ranging lions in East Africa. Clinical and Vaccine Immunology, 3(5), 554–562. Retrieved from https://www.ncbi.nlm.nih.gov/pmc/articles/PMC170405/pdf/030554.pdf

Hollings, T., Robinson, A., van Andel, M., Jewell, C., & Burgman, M. (2017). Species distribution models: A comparison of statistical approaches for livestock and disease epidemics. PLOS ONE, 12(8), e0183626. doi:10.1371/journal.pone.0183626

Kohavi, R., & John, G. H. (1997). Wrappers for feature subset selection. Artificial Intelligence, 97(1-2), 273–324. doi:10.1016/S0004-3702(97)00043-X

Kuhn, M. (2008). Building Predictive Models in *R* Using the caret Package. Journal of Statistical Software, 28(5), 1–26. doi:10.18637/jss.v028.i05

Kursa, M. B., & Rudnicki, W. R. (2010). Feature selection with the Boruta Package. Journal of Statistical Software, 36(11), 1–13. doi:10.18637/jss.v036.i11

Lachish, S., Gopalaswamy, A. M., Knowles, S. C. L., & Sheldon, B. C. (2012). Site-occupancy modelling as a novel framework for assessing test sensitivity and estimating wildlife disease prevalence from imperfect diagnostic tests. Methods in Ecology and Evolution, 3(2), 339–348. doi:10.1111/j.2041-210X.2011.00156.x

Lundberg, S., & Lee, S.-I. (2016). An unexpected unity among methods for interpreting model predictions. In NIPS 2016 Workshop on Interpretable Machine Learning in Complex Systems. Retrieved from http://arxiv.org/abs/1611.07478

Machado, G., Mendoza, M. R., & Corbellini, L. G. (2015). What variables are important in predicting bovine viral diarrhea virus? A random forest approach. Veterinary Research, 46(1), 85. doi:10.1186/s13567-015-0219-7

Marmion, M., Parviainen, M., Luoto, M., Heikkinen, R. K., & Thuiller, W. (2009). Evaluation of consensus methods in predictive species distribution modelling. Diversity and Distributions, 15(1), 59–69. doi:10.1111/j.1472-4642.2008.00491.x

Molnar, C. (2018a). iml: An R package for Interpretable Machine Learning. Journal of Open Source Software, 3(26), 786. doi:10.21105/joss.00786

Molnar, C. (2018b). Interpretable machine learning (Retrieved).

Munson, L., Terio, K. A., Kock, R., Mlengeya, T., Roelke, M. E., Dubovi, E., … Packer, C. (2008). Climate extremes promote fatal co-infections during canine distemper epidemics in African lions. PLoS ONE, 3(6), e2545. doi:10.1371/journal.pone.0002545

Nakagawa, S., & Freckleton, R. P. (2008). Missing inaction: the dangers of ignoring missing data. Trends in Ecology & Evolution, 23(11), 592–596. doi:10.1016/J.TREE.2008.06.014

Nunn, C. L., Jordán, F., McCabe, C. M., Verdolin, J. L., & Fewell, J. H. (2015). Infectious disease and group size: more than just a numbers game. Philosophical Transactions of the Royal Society of London B: Biological Sciences, 370(1669). Retrieved from http://rstb.royalsocietypublishing.org/content/370/1669/20140111.long

Ogutu, J. O., Piepho, H.-P., & Schulz-Streeck, T. (2011). A comparison of random forests, boosting and support vector machines for genomic selection. BMC Proceedings, 5(Suppl 3), S11. doi:10.1186/1753-6561-5-S3-S11

Packer, C., Altizer, S., Appel, M., Brown, E., Martenson, J., O’Brien, S. J., … Lutz, H. (1999). Viruses of the Serengeti: patterns of infection and mortality in African lions. Journal of Animal Ecology, 68(6), 1161–1178.

Penone, C., Davidson, A. D., Shoemaker, K. T., Di Marco, M., Rondinini, C., Brooks, T. M., … Costa, G. C. (2014). Imputation of missing data in life-history trait datasets: which approach performs the best? Methods in Ecology and Evolution, 5(9), 961–970. doi:10.1111/2041-210X.12232

Pollock, R. V, & Coyne, M. J. (1993). Canine parvovirus. The Veterinary Clinics of North America. Small Animal Practice, 23(3), 555–68. Retrieved from http://www.ncbi.nlm.nih.gov/pubmed/8389070

Roelke-Parker, M. E., Munson, L., Packer, C., Kock, R., Cleaveland, S., Carpenter, M., … Appel, M. J. (1996). A canine distemper virus epidemic in Serengeti lions (*Panthera leo*). Nature, 379(6564), 441–445.

Schölkopf, B., & Smola, A. J. (2002). Learning with kernels : support vector machines, regularization, optimization, and beyond. MIT Press. Retrieved from https://dl.acm.org/citation.cfm?id=559923

Shapley, L. S. (1953). Stochastic games. Proceedings of the National Academy of Sciences of the United States of America, 39(10), 1095–100. doi:10.1073/PNAS.39.10.1095

Stekhoven, D. J., & Bühlmann, P. (2012). Missforest-Non-parametric missing value imputation for mixed-type data. Bioinformatics, 28(1), 112–118. doi:10.1093/bioinformatics/btr597

Štrumbelj, E., & Kononenko, I. (2014). Explaining prediction models and individual predictions with feature contributions. Knowledge and Information Systems, 41(3), 647–665. doi:10.1007/s10115-013-0679-x

Tu, J. V. (1996). Advantages and disadvantages of using artificial neural networks versus logistic regression for predicting medical outcomes. Journal of Clinical Epidemiology, 49(11), 1225–1231. doi:10.1016/S0895-4356(96)00002-9

Viana, M., Cleaveland, S., Matthiopoulos, J., Halliday, J., Packer, C., Craft, M. E., … Lembo, T. (2015). Dynamics of a morbillivirus at the domestic–wildlife interface: Canine distemper virus in domestic dogs and lions. Proceedings of the National Academy of Sciences of the United States of America, 112(5), 1464–1469. doi:10.1073/pnas.1411623112

