## Supplementary material for "How to make more from exposure data? An integrated machine learning pipeline to predict pathogen exposure": Text S1

Text S1: Machine learning exposure risk model vignette


### Text S1: Machine learning exposure risk model vignette

###### *Nicholas M. Fountain-Jones, Gustavo Machado, Scott Carver, Craig Packer, Mariana Recamonde-Mendoza & Meggan E. Craft*

###### *4 February 2019*

This vignette will provide a worked example to running the machine learning exposure risk models outlined in Fountain-Jones et al. This particular example is for feline parvovirus exposure in the Serengeti lions. The data sets and two functions are available at github (https://github.com/gustavo-etal/Machine-learning-exposure-risk-model).

We start the pipeline by clearing the work space and loading all of the required packages.

```
library(memisc)
rm(list = ls())

library("randomForest");library("caret");library("pROC");library("ROCR");library("plyr");library("missForest")
library("gbm");library("pdp");library("ggplot2"); library("iml"); library("Boruta");library("dplyr")

# load functions
source("functions.R")
source("functionsCV.R")
```

Next we will import the parvovirus data set. There are 18 columns with one response (parvovirus seropositive or seronegative) and 17 features for 300 lions (rows).

```
# ## define column classes
data <- read.csv("parvoAgeExp.csv", row.names=1)
data$Epidemic_year <- as.factor(data$Epidemic_year)
```

### 1. Pre-processing

##### 1.1. What to do with missing data?

As you can see this data set has numerous missing data in mostly the male lions in which we did not know the pride they originated from. In this example we remove them.

```
data<-data[complete.cases(data), ] #dropping NAs
```

but we provide code for the ‘MissForest’ algorithm which may be appropriate for some data sets. If you do apply ‘Missforest’ check the normalized root mean square error (NRMSE) for continuous variables and the proportion of falsely classified (PFC) entries for categorical features to quantify how successful imputation was.

```
#set.seed(135) #this is stochastic algorithm so set seed to ensure results can be fully replicated.
#data <- missForest(data, variablewise = TRUE)
#data$OOBerror #checks error rate for missing data impuation for each variable.
# #make a dataframe again.
#data <- as.data.frame(data$ximp)
```

##### 1.2 Selecting relevant features

Not all of these features will be useful for predicting parvovirus exposure in lions, even though we did have hypotheses to why they could be important for disease exposure in general. Thus we used the following code from the ‘Boruta’ package (Kursa & Rudnicki, 2010).

```
#assign class

colnames(data)[1] <- "Class"

X = data[-which(names(data) == "Class")]
Y <- data$Class

set.seed(124)#this is another stochastic machine learning model
ImpVar <- Boruta(X, Y, doTrace = 0, maxRuns = 999)
print(ImpVar)
```

```
## Boruta performed 998 iterations in 1.290158 mins.
##  6 attributes confirmed important: Age_exposed, Epidemic_year,
## Pride_territory_size, Sex, Territory_overlap and 1 more;
##  9 attributes confirmed unimportant: Average_number_neighbours,
## Average_Ph, Despotic, FIV_PCoA1, FIV_PCoA2 and 4 more;
##  2 tentative attributes left: Avg_group_size, Habitat_quality;
```

```
#plot with x axis label vertical
plot(ImpVar, xlab = "", xaxt = "n")
lz<-lapply(1:ncol(ImpVar$ImpHistory),function(i)
  ImpVar$ImpHistory[is.finite(ImpVar$ImpHistory[,i]),i])
names(lz) <- colnames(ImpVar$ImpHistory)
Labels <- sort(sapply(lz,median))
axis(side = 1,las=2,labels = names(Labels),
     at = 1:ncol(ImpVar$ImpHistory), cex.axis = 0.7)
```

```
#only choose the variables that are relevant for subsequent models.
set <- getSelectedAttributes(ImpVar, withTentative = TRUE)#keep tentative
dataReduced <- data[which(names(data) %in% set)]
data<- cbind(data[1], dataReduced);
```

### 2. Training the models

##### 2.1 Create partitions

The next pre-processing step involves creating data partitions for the 10-fold cross validation (CV). We do this for the complete data set and for the down sampled data set where the classes (0 and 1s) are more balanced. Parvovirus prevalence is ~30% so 209/300 lions are negative. The down sampled version creates a balanced data set with equal numbers of both classes.

```
inTrain= createDataPartition(y=data$Class, p = .8, list = FALSE) #.8 of the dataset for training is reccomended 

data.train.descr <- data[inTrain,-1]
data.train.class <- data[inTrain,1]
data.test.descr <- data[-inTrain,-1]
data.test.class <- data[-inTrain,1]

# ## create folds for CV
set.seed(123)
myFolds <- createMultiFolds(y=data.train.class,k=10,times=10)

# create folds for CV, performing downsample of majority class
set.seed(130)
myFoldsDownSample <- CreateBalancedMultiFolds(y=data.train.class,k=10,times=10) 
length(unique(unlist(myFoldsDownSample)))
```

```
## [1] 116
```

```
myControl <- trainControl(## 10-fold CV)
  method = "repeatedcv",
  number = 10,
  repeats = 10,
  index = myFolds,
  savePredictions=TRUE,        
  classProbs = TRUE,
  summaryFunction = twoClassSummary,  
  allowParallel=TRUE,
  selectionFunction = "best")

#downsampled version

myControlDownSample <- trainControl(## 10-fold CV
    method = "repeatedcv",
    number = 10,
   repeats = 10,
   index = myFoldsDownSample,
   savePredictions=TRUE,        
   classProbs = TRUE,
   summaryFunction = twoClassSummary,  
   allowParallel=TRUE,
   selectionFunction = "best")
```

For large datasets you may want to register multiple cores to do the analysis in parallel. This step is skipped in this analysis.

```
#library(doParallel) 
#registerDoParallel(cores = 4)
```

##### 2.2 Running the models

Now we are ready to run the models. First we will construct a random forest (rf) model. The diagnostic plot generated assesses how the ROC (receiver operating characteristic) changes when the number of predictors (features) randomly selected to build each regression tree (mtry) is altered. In this case predictive performance is highest when only three predictors are used.

```
rf.grid <- expand.grid(.mtry=(3:6)) #number of predictors (mtry) to test.

set.seed(125) #again this is a stochastic model so set the seed to reproducibility.

rf.fit <- train(data.train.descr,data.train.class,
                method = "rf", #this is where we select which model to use for prediction (out of the 237 available)
                metric = "ROC",# Reciever opperating characteristic used to test model performance     (sensitivity/specificity)
                verbose = FALSE,
                trControl = myControl,
                tuneGrid = rf.grid, # Optimized parameter tuning is done usig the 'rf.grid' set up

                verboseIter=TRUE)

plot(rf.fit)
```

##### 2.3 Model performance

We used the following steps to calculate AUC (area under curve), specificity and sensitivity for our RF model. The mean metrics are most useful. ACC = another name for AUC, SPE = specificity, SEN: Sensitivity. For each of these metrics a score of 0.5 or below indicates a random prediction, whereas values > 0.7 (model accurately predicts negative/positive classes 70% of the time) may indicate reasonable performance for predicting exposure risk, but we urge caution as the threshold used can depend on the aims of the study.

Matthew’s correlation coefficient [MCC (Mathews, 1975)] also provides a useful metric for model performance overall. MCC values are bounded between -1 and +1 with a value of +1 represents a perfect prediction, 0 no better than random prediction and -1 indicates total disagreement between prediction and observation.

```
oof <- rf.fit$pred # check if worked
head(oof)
```

```
##       pred      obs Negative Positive rowIndex mtry     Resample
## 1 Negative Negative    0.976    0.024        6    3 Fold01.Rep01
## 2 Negative Negative    0.934    0.066       17    3 Fold01.Rep01
## 3 Negative Negative    0.644    0.356       37    3 Fold01.Rep01
## 4 Positive Positive    0.190    0.810       38    3 Fold01.Rep01
## 5 Negative Negative    0.678    0.322       54    3 Fold01.Rep01
## 6 Negative Positive    0.998    0.002       58    3 Fold01.Rep01
```

```
oof <- oof[oof$mtry==rf.fit$bestTune[,'mtry'],]# consider only the best tuned model to use

repeats<-rf.fit$control$repeats

rf.performance.cv <- EstimatePerformanceCV(oof=oof,repeats=repeats) #Matthew's correlation coefficient (MCC)
rf.performance.cv
```

```
## $mean.confusion
##               Ref Negative Ref Positive
## Pred Negative         64.8         14.1
## Pred Positive         13.2         23.9
## 
## $deviation.confusion
##               Ref Negative Ref Positive
## Pred Negative     1.549193     1.969207
## Pred Positive     1.549193     1.969207
## 
## $mean.metrics
##        ACC        SPE        SEN        MCC 
## 76.4655172 83.0769231 62.8947368  0.4626476 
## 
## $deviation.metrics
##        ACC        SPE        SEN        MCC 
## 2.50676190 1.98614531 5.18212473 0.05964692
```

```
# estimate error rates from repeated cross-validation
rf.error.cv <- EstimateErrorRateCV(oof=oof,repeats=repeats)
rf.error.cv
```

```
## $mean.positive
## [1] 37.10526
## 
## $deviation.positive
## [1] 5.182125
## 
## $mean.negative
## [1] 16.92308
## 
## $deviation.negative
## [1] 1.986145
```

```
rm(oof,repeats)

#save the model
save(rf.fit,file="Parvo Age Exposed RF.RData")
```

##### 2.4 Comparing models

We then use the same approach and syntax to compare all models (see 4.0 below for code). The model with the best predictive performance (i.e., the highest AUC, MCC, sensitivity and specificity) will be used in subsequent analysis. In this vignette, we will assume that the RF model (with no down sampling of classes) had the best overall performance and will explore this model further.

### 3. Interpreting the model

##### 3.1 Feature importance

To visualize feature performance we use the model agnostic ‘model class reliance’ method (see main text). Feature importance can be interpreted as the amount (or factor) by which model error is increased by removing this feature compared to the original model error. All plots use ggplot2 graphics and are based on code by Molnar (2018).

```
X <-data[-which(names(data) == "Class")] #load data again for the visualization
Y <- data$Class

# create the iml object

mod <-Predictor$new(rf.fit, data = X, y = Y) #create predictor object. Add your model object name here (GLM, RF, GBM or SVM)
set.seed(123)
imp <-FeatureImp$new(mod, loss = "ce", method="cartesian") #cartesian is more thorough but 'shuffle' is faster.
imp.dat<- imp$results
plot(imp)+ theme_bw()#plot results
```

##### 3.2 Partial dependency (PD) plots and centered individual conditional expectation (cICE) plots.

PD plots and cICE plots provide a valuable way to visualize what effect each predictor in the model has in shaping exposure risk. Here we plot the two most important features in our RF model (but we provide code to plot all of the relevant features). This code plots the feature effect for both classes (negative and positive). The y-axis representing the probability of parvovirus exposure. The vertical lines on the x-axis (or ‘rug plot’) shows the distribution of observations across the feature. The highlighted line in the PD plot (i.e. model average) with the other lines (the ICE lines) reflect the predictions for one observation when we vary that feature.

The age exposed cICE plots demonstrate that, whilst exposure risk of parvovirus is higher in cubs ~2 years of age, some observations have a much more subtle peak. This could be evidence for an interaction effect (i.e., risk is lower for some cubs). CICE curves are diffiult to interpret for categorical predictors so we use standard ICE plots. The resultant box plot shows parvovirus exposure risk was higher in the 1992 epidemic compared to the others.

```
#plot pd plots for the top predictors (any number appropriate for the data). Cateforical features don't plot properly

top2<- imp.dat$feature[1:2]#n = number of predictors you want to display

ice_curves <- lapply(top2, FUN = function(x) {
  cice <- partial(rf.fit, pred.var = x, center = TRUE, ice = TRUE, which.class="Positive",
                  prob = T) #note that we center values in the plotso these are centered ICE plots (cICE)
  autoplot(cice, rug = TRUE, train = dataReduced, alpha = 0.1) +
    theme_bw() +
    ylab("c-ICE")
  })

# Epidemic_year was the second most important predictor but is categorical so ICE plot is dififcult to interpret. We make a boxplot of the same data instead.

Categorical <- "Epidemic_year"
ice_curves1 <- lapply(Categorical, FUN = function(x) {
  ice <- partial(rf.fit, pred.var =  'Epidemic_year', ice = TRUE, center = FALSE, which.class="Positive",
                  prob = T)
  ggplot(ice, rug=T, train = data, aes(x=Epidemic_year, y = yhat, group = Epidemic_year)) +
    geom_boxplot()+theme_bw() 
})
#put them together
grid.arrange(grobs = c(ice_curves, ice_curves1), ncol = 2)
```

```
#for plots of a particular feature

#rf.fit %>%
  #partial(pred.var = "Age_exposed", ice=T, which.class="Positive", prob = T) %>%
  #autoplot(rug = TRUE, center =TRUE, train = data, alpha = 0.1)+theme_bw()
```

##### 3.3 Interactions using Friedman’s H index

Calculating Friedman’s H index provides a robust way to assess the importance of interactions in shaping risk across models. The interactions identified can then be visualized using PD plots.

```
set.seed(345)
mod <- Predictor$new(rf.fit, data = X, y=Y, type='prob', class='Positive') #we just want  positive class results now

interact <- Interaction$new(mod)
plot(interact)+ theme_bw()
```

This plot shows that Age\_exposed has the highest H score and is involved in the most interactions with other features in shaping parvovirus risk. We can interrogate these interactions and plot the strongest identified with the following code:

```
interact1 <- Interaction$new(mod, feature = "Age_exposed")

plot(interact1)+theme_bw()
```

```
pdp.obj <-  FeatureEffect$new(mod, feature = c("Age_exposed","Epidemic_year"), method='pdp', run=TRUE)


plot(pdp.obj)+ scale_fill_gradient(low = "white", high = "red")+ theme_bw()
```

The interaction PD plots illustrates that the risk of parvovirus exposure was much greater for cubs in the 1992 epidemic compared to the 1985 or 1977 epidemic, although we do not have data from many individuals on the landscape in 1977.

##### 3.4 Explain single predictions using Shapely values

To better understand model predictions, the last step is to use a game theory and specifically Shapely values to understand how the model is applied to individual observations (see main text for more details).

```
shapley <- Shapley$new(mod, x.interest = X[1,]) #EL BRAVO (parvovirus negative)
shapley$plot()+ theme_bw()
```

```
results <- shapley$results #for each instance you can view these results as a table
head(results, n = 7L)
```

```
##                feature      phi     phi.var                 feature.value
## 1      Habitat_quality -0.00006 0.002987188        Habitat_quality=92.652
## 2 Pride_territory_size  0.01538 0.007288137  Pride_territory_size=121.833
## 3    Territory_overlap -0.05800 0.009575758 Territory_overlap=0.787963852
## 4                  Sex -0.03982 0.008754674                         Sex=M
## 5      Yearly_rainfall -0.01754 0.001773402        Yearly_rainfall=104.58
## 6       Avg_group_size  0.00354 0.002911342              Avg_group_size=3
## 7        Epidemic_year -0.09382 0.022896169              Epidemic_year=85
```

```
sum(results$phi)# <0 indicate model prediction was negative, > 0 model prediction was postive.
```

```
## [1] -0.22068
```

The output and plot show that for the male lion named El Bravo who was parvovirus negative, the model accurately predicted as the sum of the Shapely values were negative (Shapley value = -0.22). This prediction was because, according to the model, risk was lower during the 1985 epidemic, El Bravo was male and the pride he belonged to had high overlap with other prides. However, the pride he belonged to also had a large territory size and this increased his risk of being exposed slightly (Shapley value = 0.002). We provide code to show model predictions for another individual (Lupine) who tested positive for parvovirus in the 1992 epidemic.

```
#shapley2 <- Shapley$new(mod, x.interest = X[131,]) #LUPINE (parvovirus positive)
#shapley2$plot()+ theme_bw()
```

Molnar (2018) describes other options for global and local model predictions which may be useful.

### 4. Code for the other models

Finally, below are the other models that can be tested following a same syntax as the RF model above.

```
#------------------------------------------------------------------
################# Gradient Boosting #################
#------------------------------------------------------------------
# 
# #set up GBM tuning paramters
 gbm.grid <-  expand.grid(interaction.depth = c(1,3,5,7,9),
                          n.trees = (1:30)*10,
                          shrinkage = 0.1,
                          n.minobsinnode = c(10))# will stop when is 10 onservation in terminal node

 nrow(gbm.grid)
set.seed(123)
 gbm.fit <- train(data.train.descr, data.train.class,
                  method = "gbm",
                  metric = "ROC",
                  verbose = FALSE,
                  trControl = myControl,
                  ## Now specify the exact models 
                  ## to evaludate:
                  tuneGrid = gbm.grid)
# 
 plot(gbm.fit)
save(gbm.fit,file="CDV Age Sampled GBMAll.RData")
# # estimate model performance in terms of a confusion matrix from repeated cross-validation
oof <- gbm.fit$pred
# # consider only the best tuned model
oof <- oof[intersect(which(oof$n.trees==gbm.fit$bestTune[,'n.trees']),which(oof$interaction.depth==gbm.fit$bestTune[,'interaction.depth'])),]
repeats <- gbm.fit$control$repeats
gbm.performance.cv <- EstimatePerformanceCV(oof=oof,repeats=repeats)
gbm.performance.cv
# # estimate error rates from repeated cross-validation
gbm.error.cv <- EstimateErrorRateCV(oof=oof,repeats=repeats)
gbm.error.cv 

#------------------------------------------------------------------
################# Support vector machine #################
#------------------------------------------------------------------
# 
# 
set.seed(123)
svm.fit <- train(Class ~ .,data=cbind(Class=data.train.class,data.train.descr),
                 method = "svmRadial",
                 tuneLength = 9,
                 metric="ROC",
                 trControl = myControl)
plot(svm.fit)
# 
# # estimate model performance in terms of a confusion matrix from repeated cross-validation
oof <- svm.fit$pred
# # consider only the best tuned model
oof <- oof[intersect(which(oof$sigma==svm.fit$bestTune[,'sigma']),which(oof$C==svm.fit$bestTune[,'C'])),]
repeats <- svm.fit$control$repeats
svm.performance.cv <- EstimatePerformanceCV(oof=oof,repeats=repeats)
svm.performance.cv 
# # estimate error rates from repeated cross-validation
svm.error.cv <- EstimateErrorRateCV(oof=oof,repeats=repeats)
rm(oof,repeats)
# save the model
save(svm.fit, file="CDVAgeSampledSVM_down.RData")

#------------------------------------------------------------------
################# GLM - Logistic regression #################
#------------------------------------------------------------------

#for GLM as has problems with multi-level factors with some with sparse data in some levels, so the following step needs to be added to the pre-processing steps prior to analysis.
data <- select (data,-c(Epidemic_year))

glm.fit <- train(data.train.descr,data.train.class,
                 method = "glm",
                 metric="ROC", # will use accuracy instead
                 family="binomial",
                 trControl=myControlDownSample)


# estimate model performance in terms of a confusion matrix from repeated cross-validation
oof <- glm.fit$pred
# consider only the best tuned model
oof <- oof[which(oof$parameter==glm.fit$bestTune[,'parameter']),]
repeats<-glm.fit$control$repeats
glm.performance.cv <- EstimatePerformanceCV(oof=oof,repeats=repeats)
glm.performance.cv 

# estimate error rates from repeated cross-validation
glm.error.cv <- EstimateErrorRateCV(oof=oof,repeats=repeats)
rm(oof,repeats)


save(glm.fit,glm.performance.cv,glm.error.cv,file="model_glm_fullfeatset_Atb.RData")
```

### References

Kursa, M. B., & Rudnicki, W. R. (2010). Feature selection with the Boruta Package. Journal of Statistical Software, 36(11), 1–13. doi:10.18637/jss.v036.i11

Matthews, B. W. (1975). “Comparison of the predicted and observed secondary structure of T4 phage lysozyme”. Biochimica et Biophysica Acta (BBA) - Protein Structure. 405 (2): 442–451

Molnar, C. (2018). Iml: An R package for Interpretable Machine Learning. Journal of Open Source Software, 3(26), 786. doi:10.21105/joss.00786
