## Supplemental files for "How to make more from exposure data? An integrated machine learning pipeline to predict pathogen exposure"

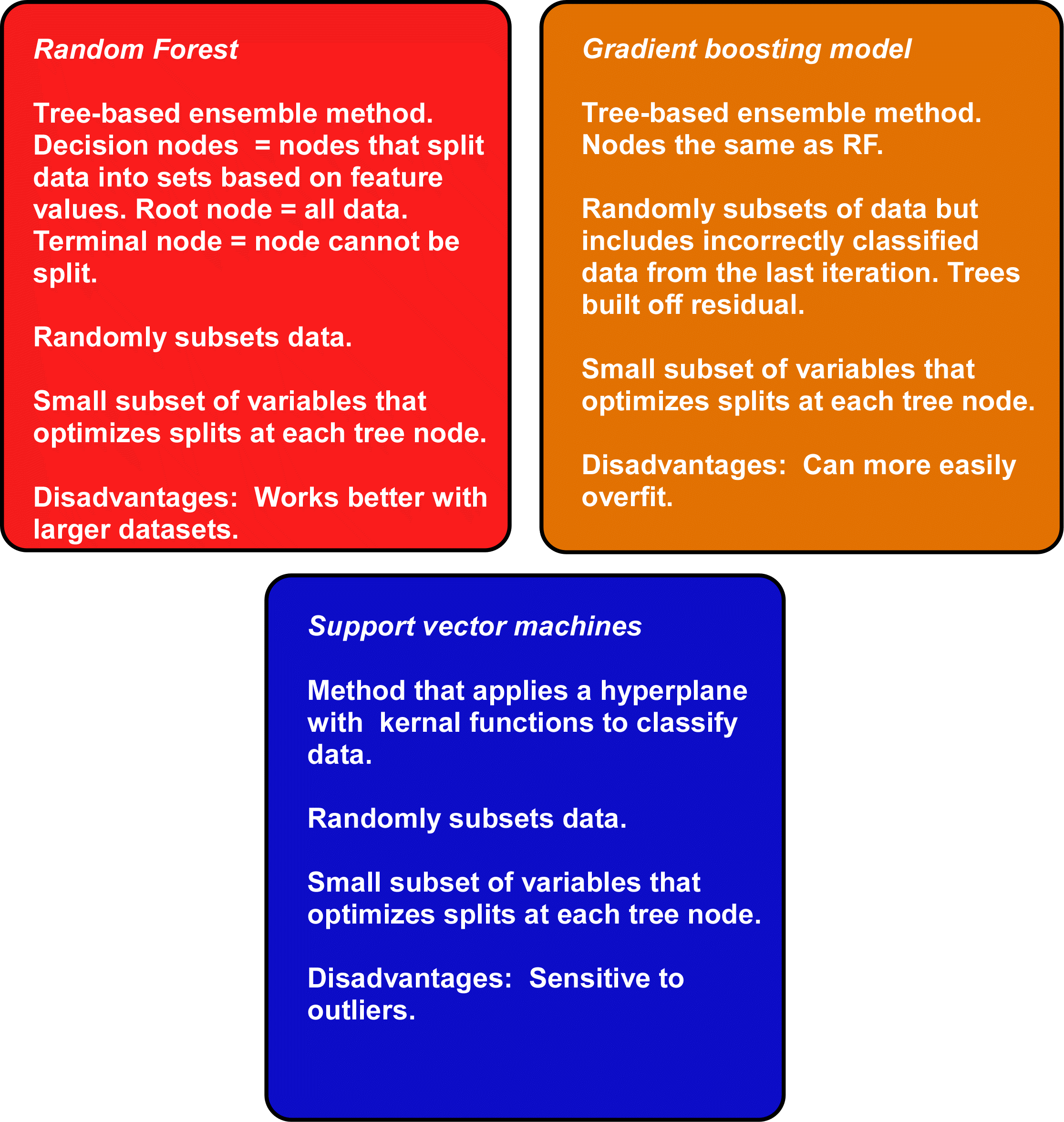

**Fig. S1:** Summary of machine learning methods we compare in our pipeline.

**Text S2: Feature selection details**

We selected features that we thought were likely to be important for pathogen exposure including variables capturing individual variability, and other pride characteristics including environmental variables (see Table S1 for measurement details and hypotheses). Because the exposure risk of CDV and parvovirus may be affected by feline immunodeficiency virus (FIV_Ple_) subtype infections (Troyer et al., 2011), we calculated two FIV_Ple_ predictors. Prevalence of FIV_Ple_ was 100% in the SLP population, although subtype prevalence varied between three subtypes (A, B & C) (see Troyer et al., 2011 for FIV relevent methods). Rather than adding infection status of each subtype, and for each subtype co-infection combination (e.g., individuals co-infected by FIV_Ple_ subtypes A/B, B/C, A/C, or A/B/C), we summarised infection information using principal coordinate analysis (PCoA). This reduced the FIV_Ple_ data to two PCoA axes that accounted for 100% of variation in the dataset; each axis was included as a variable in each model (see Fig. S2). PCoA was conducted in PRIMER- E PERMANOVA+ software (Anderson, Gorley, & Clarke, 2008). Age was included as either ‘age sampled’ or ‘age exposed’ in each model.

As both of the pathogens were epidemic in our lion population (Fig. 1), we calibrated all features to likely epidemic year and calculated ‘age exposed’ ‘time since epidemic’ and ‘year exposed’. We focussed on the interactions including ‘epidemic year’ to evaluate if the ecological context underlying exposure risk differed with each year. In total, the CDV models included data for 184 individuals on the landscape in 1981 and 1994 respectively, whereas the parvovirus models included 285 individuals on the landscape in 1985 and 1992 respectively. For CDV, only 2 individuals were potentially exposed to the 1977 epidemic, so these were removed from the analysis. In both models, individuals that were not on the landscape during either epidemic were excluded. However, to account for uncertainty in the epidemic year estimates, if an individual was born that epidemic year we included them even though it is possible that their antibodies were maternally derived. See Table S1 for a summary of what features were included in each analysis.

Most missing data was concentrated in males that were not born in the SLP population and had arrived from elsewhere and ‘missForests’ predictions often poorly predicted this data. Thus, we made the decision to exclude missing data from all analyses.

**Table S1.** Details of the individual, pride and environmental level features used in each model.

| **Feature** | **Type** | **Measurement details** | **Hypothesis?** | **Data** | **Model used** |
| --- | --- | --- | --- | --- | --- |
| Sex | Individual | Sex determined during sample collection. | Males could be more likely to be exposed due to physiological stress. | SLP data | All |
| Age sampled | Individual  (Temporal) | Age at time sampled | Older individuals could be more likely exposed. | SLP data | U |
| Age exposed | Individual  (Temporal) | Age at time potentially exposed | Older individuals could be more likely exposed. | SLP data | CEp |
| Epidemic year | Individual  (Temporal) | Year individual was likely to be exposed. | Some epidemic years may have higher risk than others. | SLP data | CEp |
| Time since epidemic | Individual  (Temporal) | The number of years between likely exposure year and the year sampled. | Detection of exposure events could decrease with time since epidemic due to waning titer levels. | SLP data | CEp |
| Number of immigration events  (# immigrations) | Individual | Number of prides an individual has successfully immigrated into prior to sampling. | Individuals that have immigrated multiple times may be exposed. | SLP data | **All** |
| Pride, coalition or nomad?  (Pride or nomad?) | Individual | Was the individual pride-living or nomadic in the previous two years prior to exposure of infection (binary). | Pride-living or males dominating multiple prides in coalitions may be more likely higher exposure risk due to higher contact rates. | SLP data | **All** |
| FIV_Ple_ B/C  (FIV (B/C)) | Individual | PCoA axis 1 scores that distinguish between individuals infected or co-infected with FIV_Ple_ B/C. | Individuals are infected or co-infected by FIV_Ple_ B or C may have higher risk of exposure to other pathogens due to T-cell depletion. | (Troyer et al., 2011) | All |
| FIV_Ple_ A  (FIV (A)) | Individual | PCoA axis scores that distinguish between individuals infected or co-infected with FIV_Ple_ A. | Individuals are infected or co-infected by FIV_Ple_ A may have higher risk of exposure to other pathogens due to T-cell depletion. | (Troyer et al., 2011) | All |
| Group size | Pride | Average number of individuals in pride two years prior sample collection. | Individuals in larger groups may be more likely to be exposed to more pathogens due to higher densities. | SLP data | **All** |
| Group behaviour | Pride | Was the pride the individual was in despotic? | Despotic prides have more interactions thus are more likely to be exposed to a greater number of directly transmitted diseases. | SLP data | **All** |
| Territory size | Pride | Based on 75% kernal. | Prides that have a larger territory may be exposed to more pathogens. | SLP data | **All** |
| Territory overlap | Pride | What percentage of prides territory overlapped with other prides. | Individuals in prides that have more overlap may have higher risk of being exposed to a pathogen. | SLP data | **All** |
| Habitat quality* | Pride | Pride habitat quality score. | Better territories are closer to rivers. Better territories require more aggressive interactions to defend which could increase exposure risk. | (Mosser, Fryxell, Eberly, & Packer, 2009) | **All** |
| Number of neighbours  (Neighbour #) | Pride | Number of individuals in neighbouring prides. Neighbouring prides had territory overlap. | Individuals in prides in high density areas will have more contacts thus are have higher exposure risk | SLP data | **All** |
| Average monthly rainfall* | Environmental | Average monthly rainfall experienced in each pride’s territory based on weather stations in the plains and woodlands. | Higher rainfall could impact prey density thus changing lion behaviour and increasing exposure risk. | (Sinclair et al., 2013) | **All** |
| Vegetation* | Environmental | Average vegetation cover across the prides territory. | Increased vegetation cover could impact prey density thus changing lion behaviour and increasing exposure risk. | (Reed, Anderson, Dempewolf, Metzger, & Serneels, 2009) | **All** |

U: Uncalibrated model, CEp: Calibrated epidemic pathogen, CEn: Calibrated endemic pathogen, All: feature used for all models (bold indicates that the feature was calibrated). * indicates that we averaged this feature across the pride’s territory.

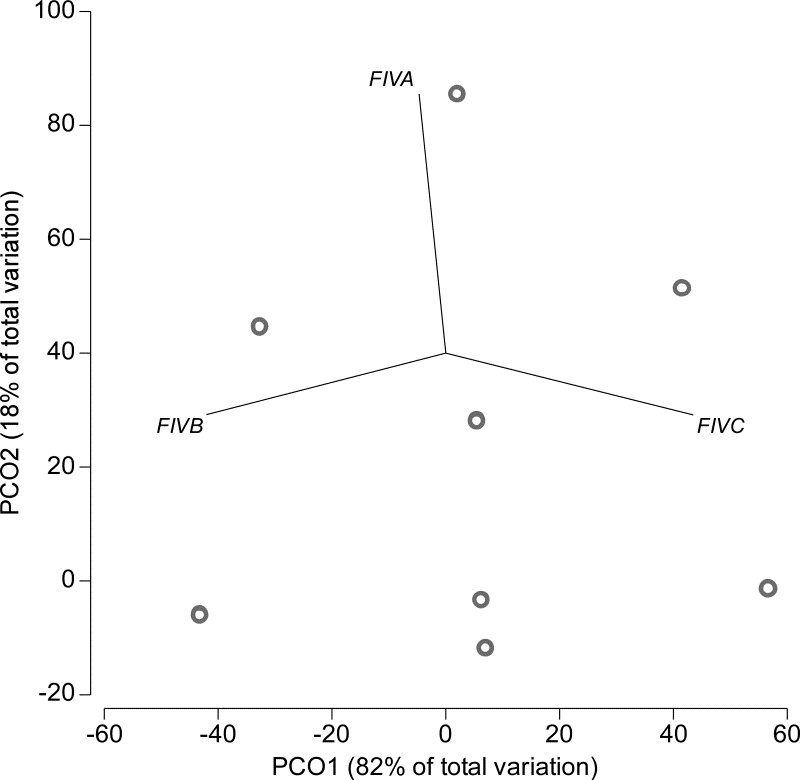

**Fig. S2:** Ordination of FIV_Ple_ infection data. For this analysis, infection data was transformed into a Jaccard similarity matrix prior to analysis. Grey circles reflect individual FIV_Ple_ subtype infections with many circles overlapping (i.e., many individuals are just infected by just FIV_Ple_ B, therefore occuping the same postion in the bottom left of the ordination). PCoA 1 (“PCO1”) accounted for 82% of the variation with negative axis values associated with individuals infected by FIV_Ple_ B, values around 0 were associated with individuals co-infected by FIV_Ple_ B and C. PCoA 2 (“PCO2”) explained 18% of the variation sepparated individuals infected by just FIV_Ple_ B (negative values), FIV_Ple_ A (values > 80) or coinfected by both (values 40-60).

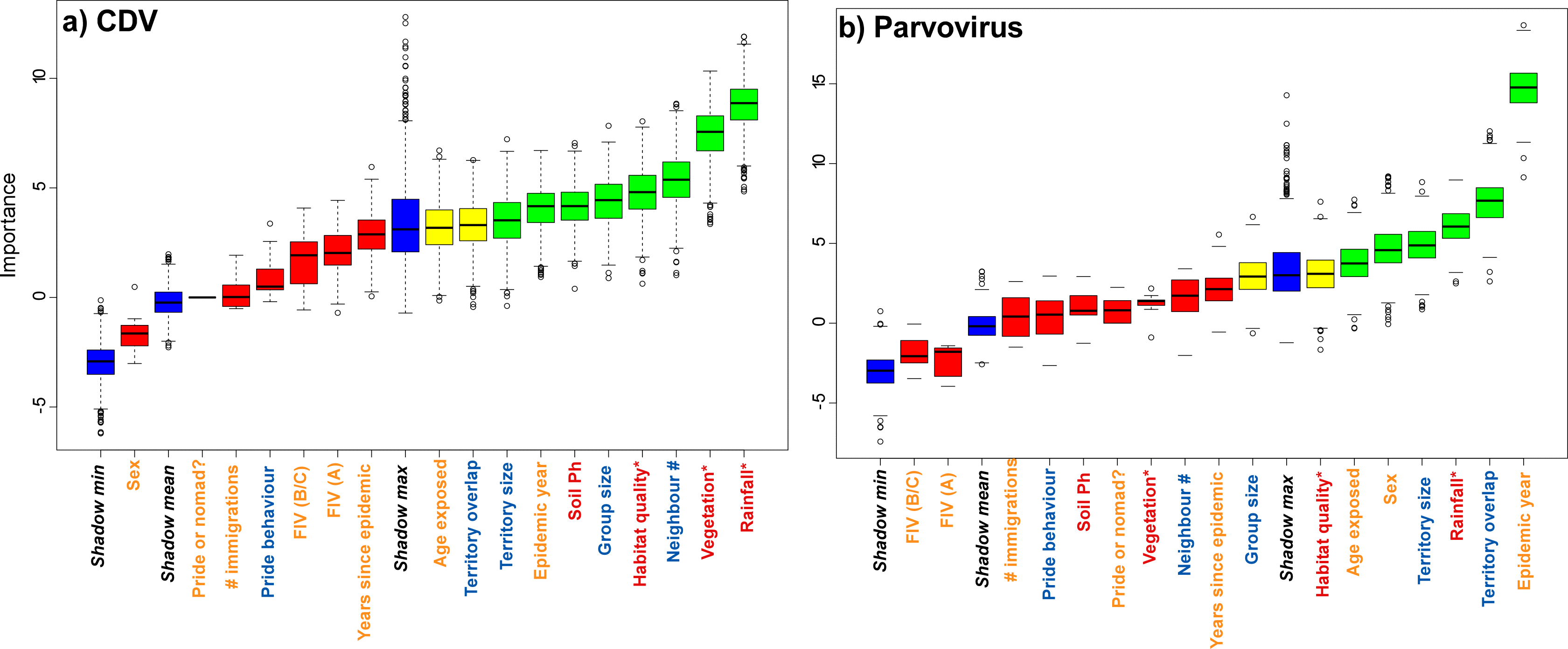

**Fig. S3:** Boruta boxplots for a) CDV and b) parvovirus calibrated models. Blue boxplots correspond to minimal, average and maximum Z score of shadow features (see Kursa & Rudnicki, 2010). Red, yellow and green boxplots represent Z scores of rejected, tentative and confirmed features respectively. For both models tentative features were included.

**Table S2:** AUC, sensitivity and specificity for the GBM and SVM down sampled/non-down sampled calibrated models as well as the RF (down-sampled) model which had less predictive performance compared to RF (without down-sampling). See Table 1 for results from the best predictive model.

| Model | AUC | Specificity | Sensitivity |
| --- | --- | --- | --- |
| **CDV** |  |  |  |
| *GBM (ND)* | 82.3 | 60 | 92.3 |
| *GBM (D)* | NA | NA | NA |
| *RF (D)* | 70.53 | 72.5 | 70.41 |
| *SVM (ND)* | 80.70 | 49.38 | 93.72 |
| *SVM (D)* | 70.63 | 66.25 | 70.87 |
| **PARVOVIRUS** |  |  |  |
| *GBM (ND)* | 72.5 | 85.38 | 46.05 |
| *GBM (D)* | 71.6 | 72.20 | 64.34 |
| *RF (D)* | 65.48 | 64.82 | 73.16 |
| *SVM (ND)* | 69.30 | 72.20 | 43.05 |
| *SVM (D)* | 71.64 | 72.46 | 62.11 |

D: Down-sampled model, ND: Model with no down-sampling.

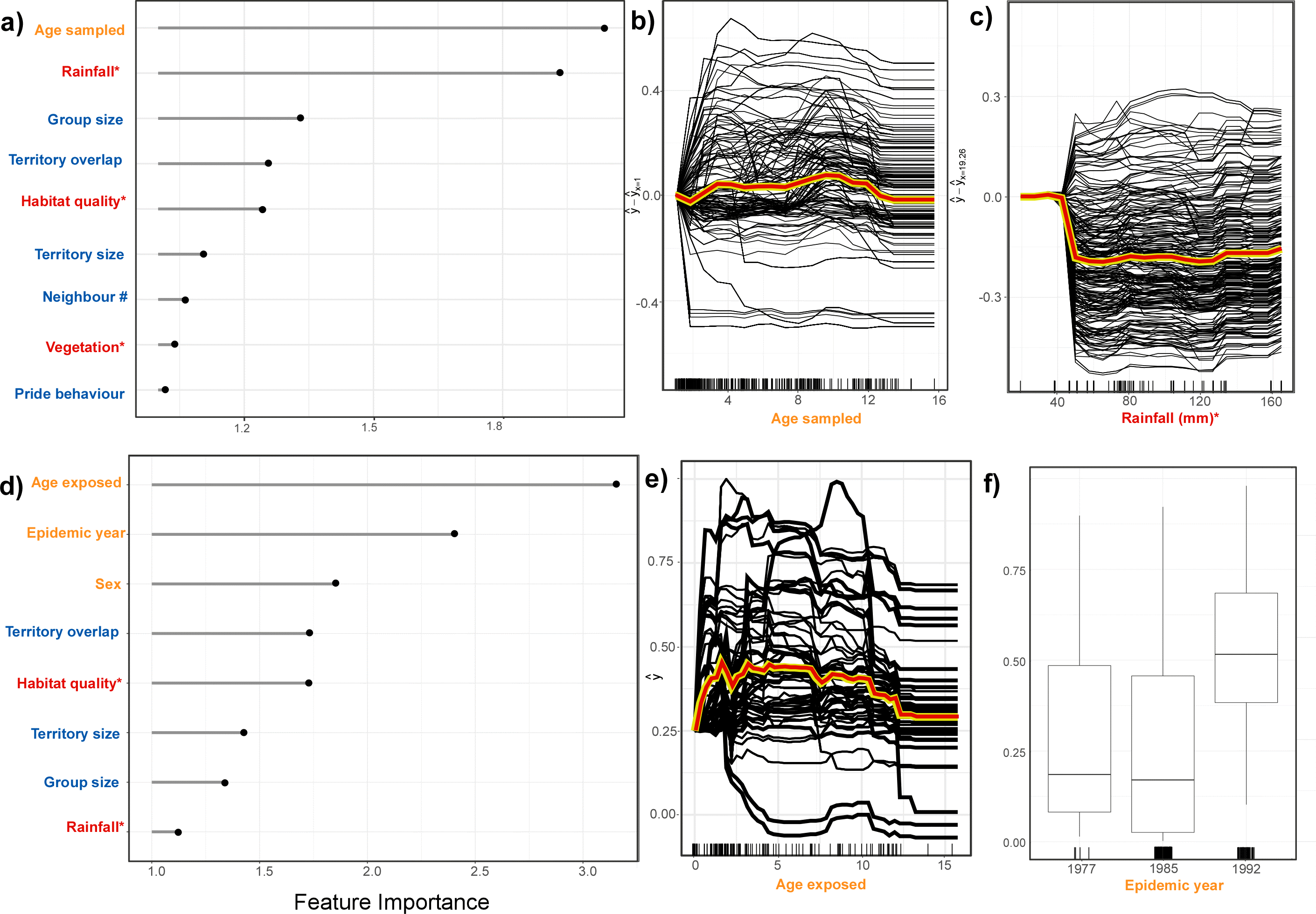

**Fig. S4:** Plots showing the differences in model predictions and the features that contribute to parvovirus exposure risk in the Serengeti lions for the uncalibrated models (a-c) compared to the calibrated models (d-f). Age sampled followed by rainfall are the most important features associated with parvovirus exposure risk in the uncalibrated models (a). In contrast, age exposed followed by epidemic year were the most important features in the calibrated model (d). CICE plots show that the relationship with exposure risk an age exposed differs to age sampled (b: uncalibrated model, e: calibrated model). (f) ICE plot of exposure risk across epidemic years (categorical). Feature name colours reflect feature type (orange = individual, blue = pride-level, red = environmental) and * indicates that the variable was averaged across the prides territory. Tick marks on the x-axes (b, c, e & f) show the distribution of data. Yellow/red line indicates the average value across all individuals. The y axes reflect probability of being positive for parvovirus.

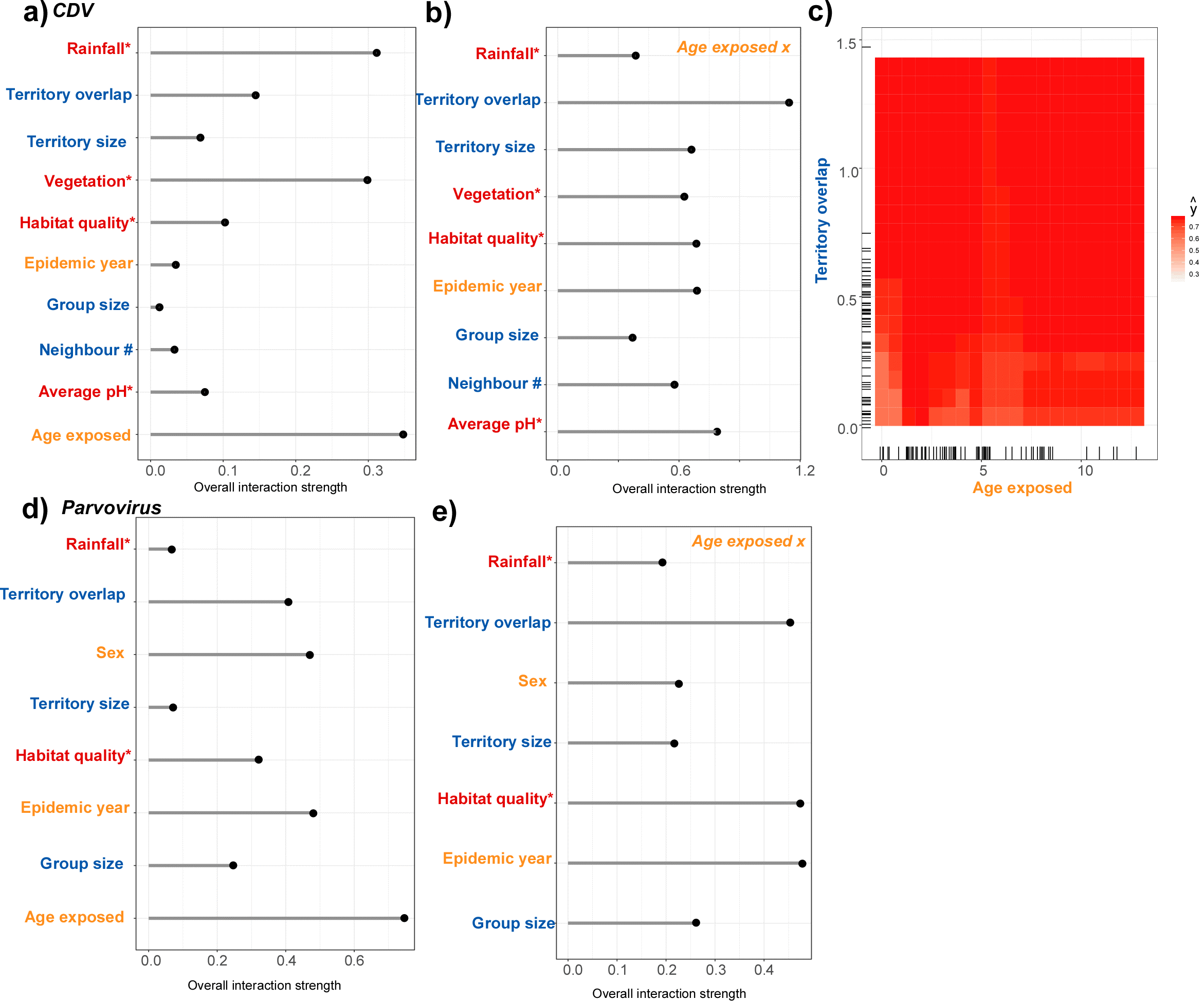

**Fig S5:** Interaction plots from the calibrated models for (a-c) CDV and (d, e) parvovirus. (a & d) show which features most interact with others to predict exposure, (b & e) show the strength of the interactions for age exposed with all other features, and (c) illustrates a interaction as a heatmap with dark red indicating relatively high risk whereas light red represents realtively lower risk. Lines on the axes (rug plots) show the distribution of each feature. The top interacting feature for parvovirus is displayed in Fig. 4 in the main text.

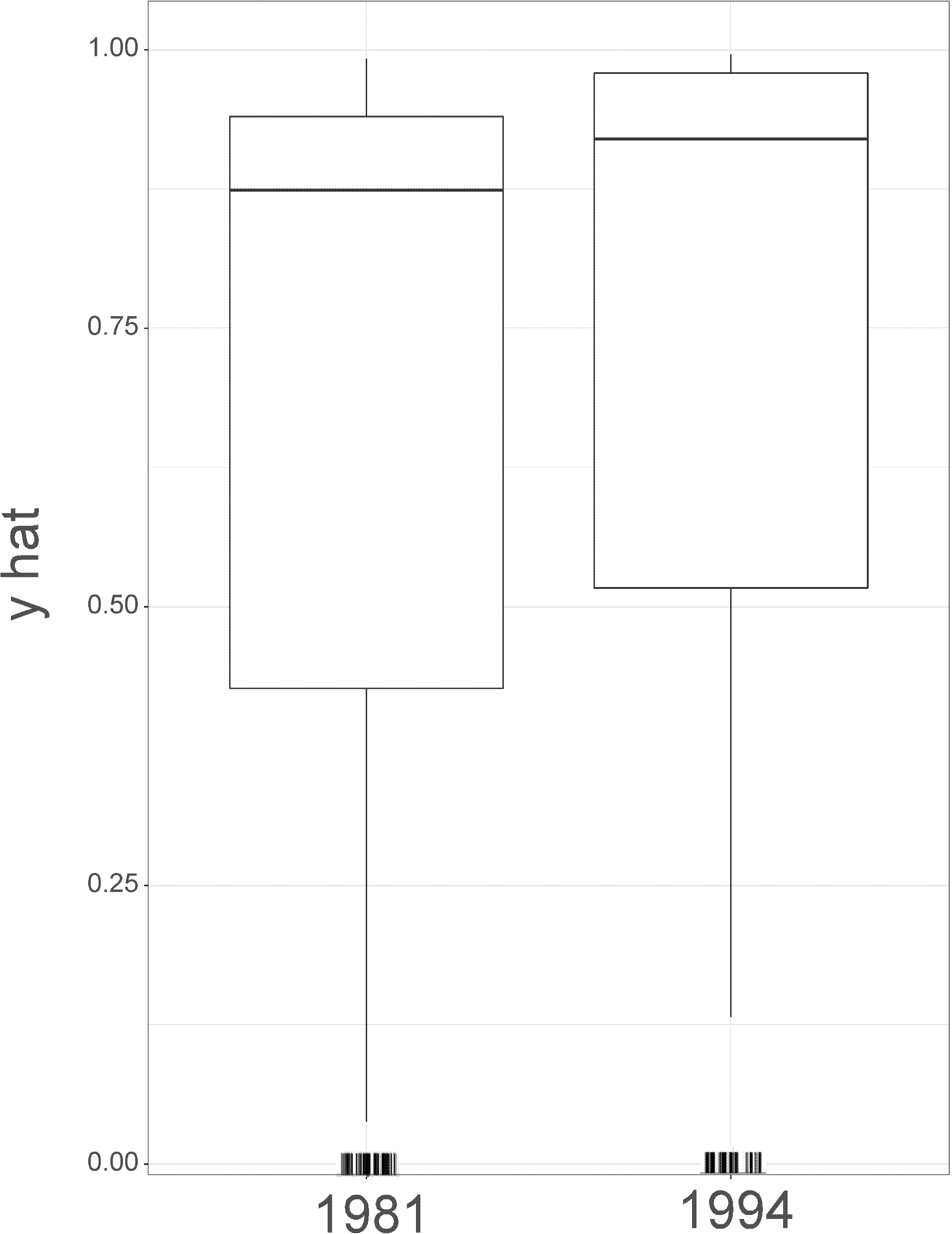

**Fig S6:** Partial dependency plot of the callibrated CDV model showing exposure risk over both epidemic years.

#### **Text S3: Game theory to understand model output**

The game theory approach helped us interpret how each feature was shaping exposure risk of each pathogen on an individual level (Fig. S5). For example, the individual ‘CS82’ from the Campsite pride (Fig. S8) tested positive to CDV (she was likely exposed in the 1994 epidemic) and, using cross validation, the model predicted this individual would be positive (Fig. S7a). The features with the highest Shapely values (and thus the drivers of this prediction) were rainfall and territory size. Summarizing, ‘CS82’ was likely exposed as she was on the landscape during an epidemic year with low rainfall in the woodlands (Fig. S8) and she belonged to a pride with relatively small territory size (Fig. S7a). Conversely, CSW from the same pride (but sampled 9 years earlier) tested negative to CDV as she was on the landscape as a cub (< 1 y.o.) during an epidemic year with relatively high rainfall and belonged to a pride with relatively high vegetative cover (Fig. S7b). ‘Askival’ tested positive to parvovirus and was on the landscape during the 1992 epidemic (Fig. S7c) in the Plains pride (Fig. S8) that had relatively low territory overlap (Fig. 5a). ‘Askival’ was a male and this made him, according to the model, slightly less likely to be exposed (negative Shapely values), however, this was outweighed by the other features with relatively high positive Shapely values (Fig. S7c). ‘7SD’ in contrast, was tested negative for parvovirus as he was on the landscape during a the lower risk 1985 epidemic and was a member of a pride K2 (Fig. S7) with a relatively high overlap and a large territory size (Fig. S7d).

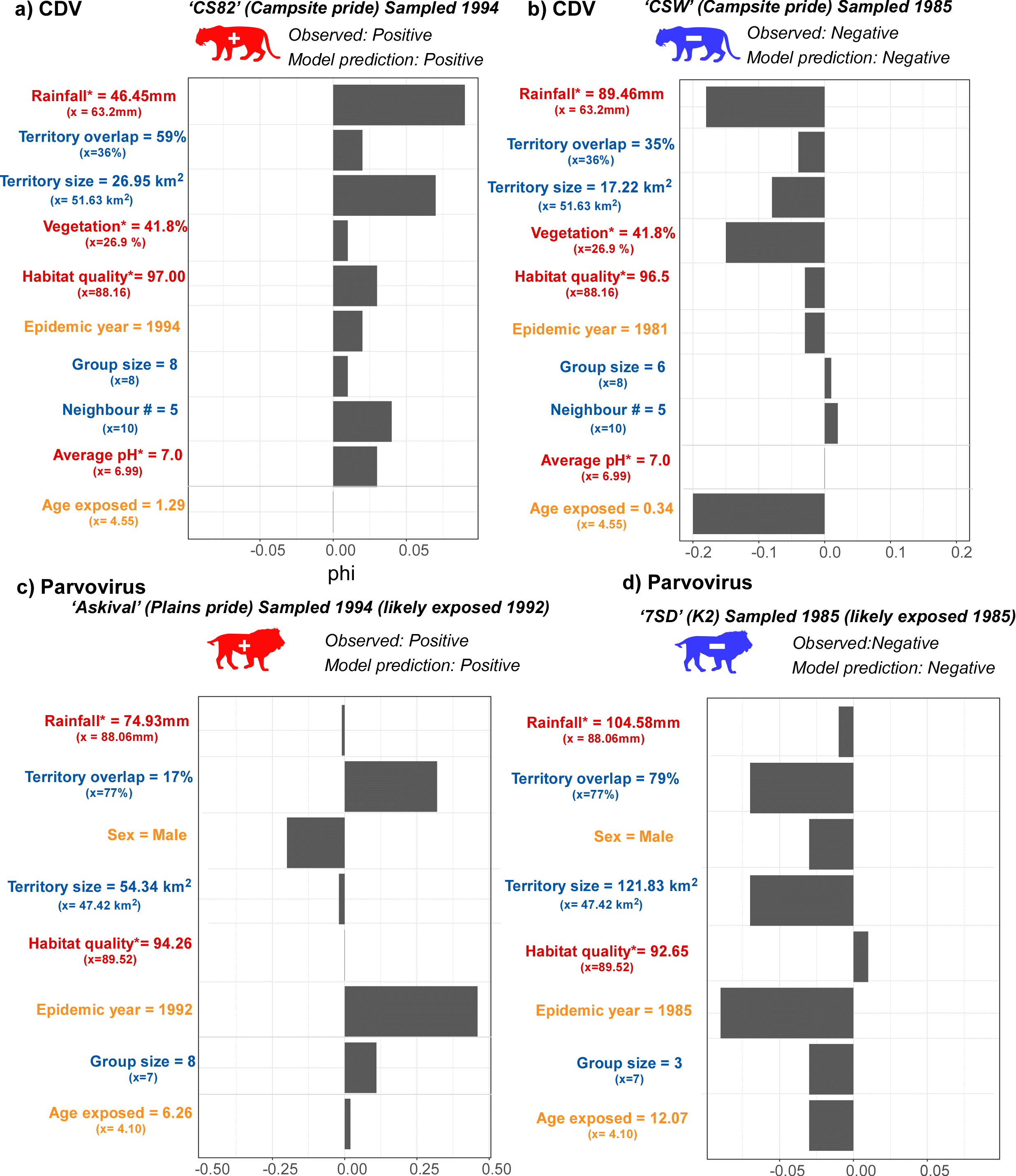

**Fig. S7:** Feature value contributions for each respective exposure calibrated model based on Shapely values for four individual lions. a) CS82’ that tested positive for CDV in 1994; b) ‘CSW’ that tested negative for CDV in 1985; c) ‘Askival’ that tested positive for parvovirus in 1994; and d) ‘7SD’ who tested negative for parvovirus in 1985. Positive Shapely values indicate that this variable contributed positively to exposure risk, whereas negative values indicate that that predictor lowered risk of exposure for that individual. The number next to each predictor name indicates the observed value of that predictor for that individual and the mean (x) of the individuals in the model is included underneath.

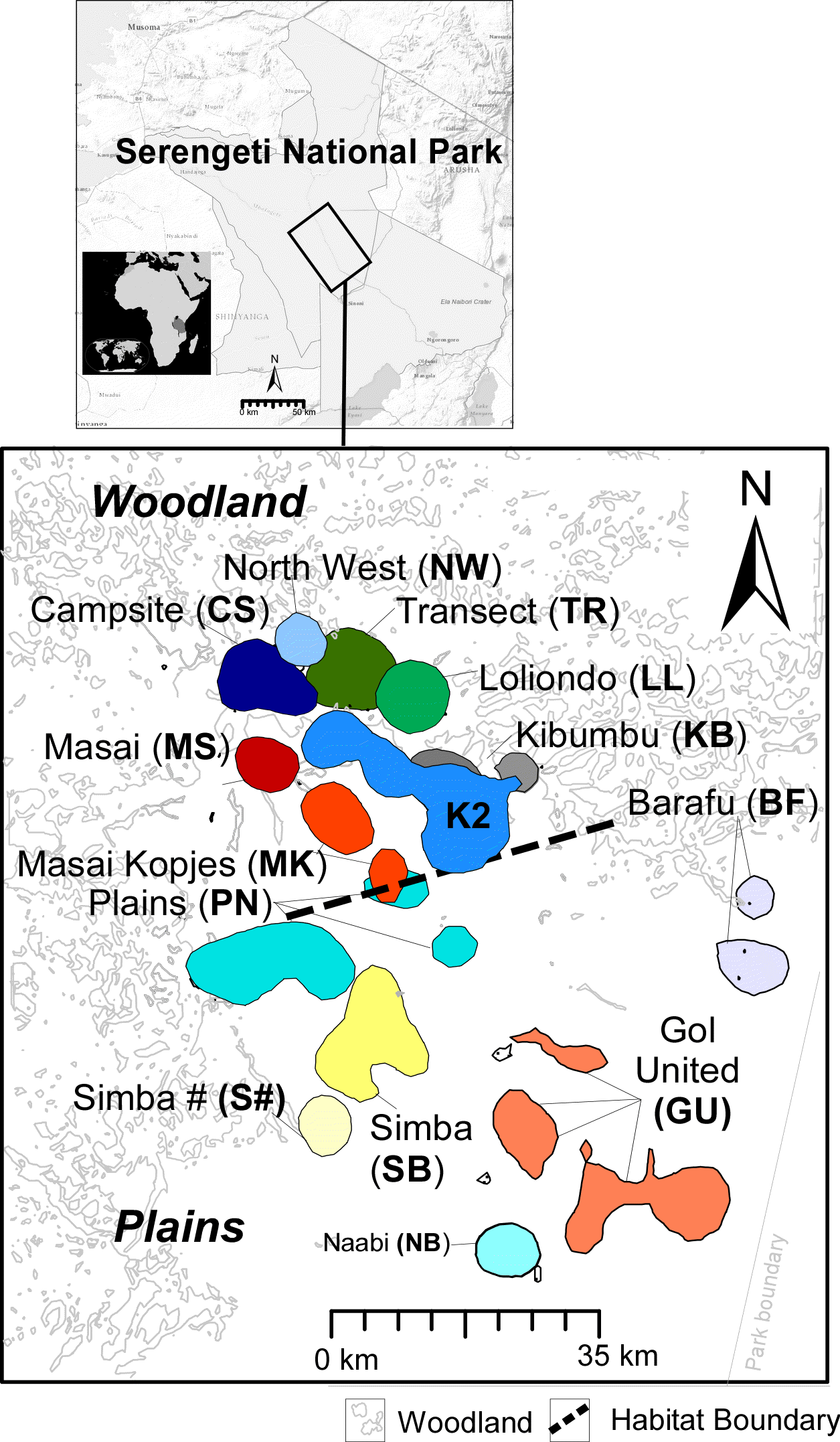

**Fig. S8:** Map of the Serengeti lion pride territories based on 1986-1987 70% territory kernel. Colours represent different pride territories. Modified from (Fountain-Jones et al., 2017).

**References**

Anderson, M. J., Gorley, R. N., & Clarke, K. R. (2008). *PERMANOVA+ for PRIMER: Guide to Software and Statistical Methods.* *2008*. Plymouth, U.K.: PRIMER-E.

Fountain-Jones, N. M., Packer, C., Troyer, J. L., VanderWaal, K., Robinson, S., Jacquot, M., & Craft, M. E. (2017). Linking social and spatial networks to viral community phylogenetics reveals subtype-specific transmission dynamics in African lions. *Journal of Animal Ecology*, *86*(6), 1469–1482. doi:10.1111/1365-2656.12751

Kursa, M. B., & Rudnicki, W. R. (2010). Feature selection with the Boruta Package. *Journal of Statistical Software*, *36*(11), 1–13. doi:10.18637/jss.v036.i11

Mosser, A., Fryxell, J. M., Eberly, L., & Packer, C. (2009). Serengeti real estate: density vs. fitness-based indicators of lion habitat quality. *Ecology Letters*, *12*(10), 1050–1060. doi:10.1111/j.1461-0248.2009.01359.x

Reed, D. N., Anderson, T. M., Dempewolf, J., Metzger, K., & Serneels, S. (2009). The spatial distribution of vegetation types in the Serengeti ecosystem: the influence of rainfall and topographic relief on vegetation patch characteristics. *Journal of Biogeography*, *36*(4), 770–782. doi:10.1111/j.1365-2699.2008.02017.x

Sinclair, A. R. E., Metzger, K. L., Fryxell, J. M., Packer, C., Byrom, A. E., Craft, M. E., … Mduma, S. A. R. (2013). Asynchronous food-web pathways could buffer the response of Serengeti predators to El Niño Southern Oscillation. *Ecology*, *94*(5), 1123–1130. doi:10.1890/12-0428.1

Troyer, J. L., Roelke, M. E., Jespersen, J. M., Baggett, N., Buckley-Beason, V., MacNulty, D., … O’Brien, S. J. (2011). FIV diversity: FIV _Ple_ subtype composition may influence disease outcome in African lions. *Veterinary Immunology and Immunopathology*, *143*(3–4), 338–346.
